# Simulation-Guided Exploration of PAINT Parameter Space for Accurate Molecular Quantification

**DOI:** 10.1101/2025.08.04.668137

**Authors:** Wei Shan Tan, Arthur M. de Jong, Menno W. J. Prins

**Affiliations:** Department of Biomedical Engineering, Eindhoven University of Technology, Eindhoven, 5612 AZ, The Netherlands; Institute for Complex Molecular Systems (ICMS), Eindhoven University of Technology, Eindhoven, 5612 AZ, The Netherlands; Department of Applied Physics and Science Education, Eindhoven University of Technology, Eindhoven, 5612 AZ, The Netherlands; Helia Biomonitoring, Eindhoven, 5612 AR, The Netherlands

## Abstract

Molecular quantification using Point Accumulation for Imaging in Nanoscale Topography (PAINT) depends critically on probe kinetics, imaging conditions, and surface molecular properties. This study presents a simulation-guided framework to systematically explore the PAINT parameter space and identify conditions that enable molecular quantification with high accuracy. Detection thresholds for key PAINT outputs, namely point spread function density, localization cloud density, and binding event density, are defined to ensure ≥ 90% accuracy in density estimates and statistical interpretability of spatial distributions. A neural network surrogate model, trained on Monte Carlo simulations, is used to perform Sobol sensitivity analysis, revealing that probe kinetics and concentration are the dominant contributors to output variability. The model also enables rapid mapping of viable parameter regimes and shows that interpretable quantification of spatial distributions in high-density, clustered systems requires either a priori knowledge of the molecular architecture or improved spatial resolution. Overall, this framework provides quantitative guidance for optimizing PAINT experiments and supports the rational design of non-DNA-based, PAINT-compatible probes, thereby expanding the applicability of PAINT to a broader range of molecular systems.

## 1 Introduction

Single-molecule localization microscopy (SMLM) comprises a family of imaging techniques that enable visualization and quantification of surface-bound biomolecules with single-molecule resolution. ^1^These techniques have provided critical insights into how the spatial organization of molecules relates to their functional roles. By temporally separating the fluorescence emissions of individual fluorophores and precisely localizing their spatial positions, SMLM surpasses the diffraction limit of conventional optical microscopy. This temporal separation, or “blinking” behavior, can be controlled through various mechanisms: by tuning the imaging buffer, as in Stochastic Optical Reconstruction Microscopy (STORM);^2^by photoactivation of fluorophores, as in Photoactivated Localization Microscopy (PALM);^3,4^or by exploiting transient binding interactions between dye-labeled probes and target molecules, as in DNA Point Accumulation for Imaging in Nanoscale Topography (DNA-PAINT).^5,6^

Among these techniques, DNA-PAINT offers several advantages, including well-defined binding kinetics and relatively simple instrumental requirements. These features make it a promising candidate for high-throughput molecular quantification. DNA-PAINT has been successfully applied to both synthetic biofunctionalized surfaces (e.g., DNA origami, nanoparticles) and biological substrates, and has been used to image diverse biomolecules including nucleic acids, proteins, and lipids.^5,7–11^Despite this versatility, molecular quantification using DNA-PAINT has largely been limited to low-density systems, those in which the intermolecular distances exceed the spatial resolution limit of the imaging method.

To extend DNA-PAINT to higher molecular densities, the quantitative PAINT (qPAINT) approach was developed, which infers molecular counts from hybridization kinetics in resolution-limited areas.^11,12^However, qPAINT requires restrictive imaging conditions such that the occurrence of multiple binding events at any given time within the same diffraction-limited region is avoided. This constraint limits its scalability for high-throughput analysis and precludes detailed assessment of spatial molecular distributions. An alternative, Resolution Enhancement by Sequential Imaging (RESI), overcomes this limitation through DNA barcoding, but suffers from poor scalability with increasing molecular density.^13^In our previous work, we introduced a simulation-based analysis framework that enables both molecular quantification and spatial distribution analysis in high-density regimes, while maintaining relatively short acquisition times.^14^

Taken together, these methodologies highlight a critical trade-off: to achieve accurate molecular quantification with high throughput, one must carefully balance probe binding kinetics and imaging parameters for the surface molecular properties under study. While DNA-PAINT benefits from predictable DNA hybridization kinetics, it also depends on tagging non-DNA targets with labeling probes, such as antibodies or nanobodies, conjugated to a DNA docking strand. These labels can introduce errors and perturb the molecular environment, potentially compromising measurement accuracy.^15^This underscores the need for novel, non-DNA-based probes that exhibit PAINT-compatible binding kinetics. To inform the development of such probes and to guide experimental design, a deeper understanding of how PAINT experimental parameters influence quantification outcomes is essential.

In this article, we present a simulation-driven strategy to systematically explore the PAINT parameter space with a focus on achieving accurate molecular quantification. We employ the ThunderSTORM simulation engine alongside custom Monte Carlo models to delineate the quantification limits for molecular density and distribution. Furthermore, we develop a neural network surrogate model to evaluate the sensitivity of PAINT outputs to individual experimental parameters. The trained model reveals optimal parameter combinations that support molecular quantification with up to 90% accuracy, and provides actionable insight into the kinetic requirements for next-generation PAINT probes.

## 2 Result and Discussion

### 2.1 Spatial Molecular Quantification Limit

Figure 1 outlines the core research questions addressed in this work. This study investigates how probe kinetic constants, imaging parameters, and the properties of the surface molecular system influence the accuracy of molecular quantification in Point Accumulation for Imaging in Nanoscale Topography (PAINT), a form of single-molecule localization microscopy (SMLM). The core principle of SMLM lies in the temporal separation of fluorescent emissions, ensuring that point spread functions (PSFs) do not overlap within any given frame. This separation is essential for the precise localization of each PSF. Consequently, there is an inherent upper limit to the PSF density, denoted as 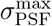, that can be tolerated in a single measurement frame. To quantify this limit, simulated PAINT datasets were generated using the ThunderSTORM simulation engine,^16^consisting of 100 frames with varying values of *σ*_PSF_ (PSF width ranging from 100 to 200 nm), serving as ground truth (see Supplementary Table S1 for simulation details). These simulated datasets were then analyzed using ThunderSTORM to obtain the localized PSF positions, which were compared against the ground truth.

**Figure 1.**
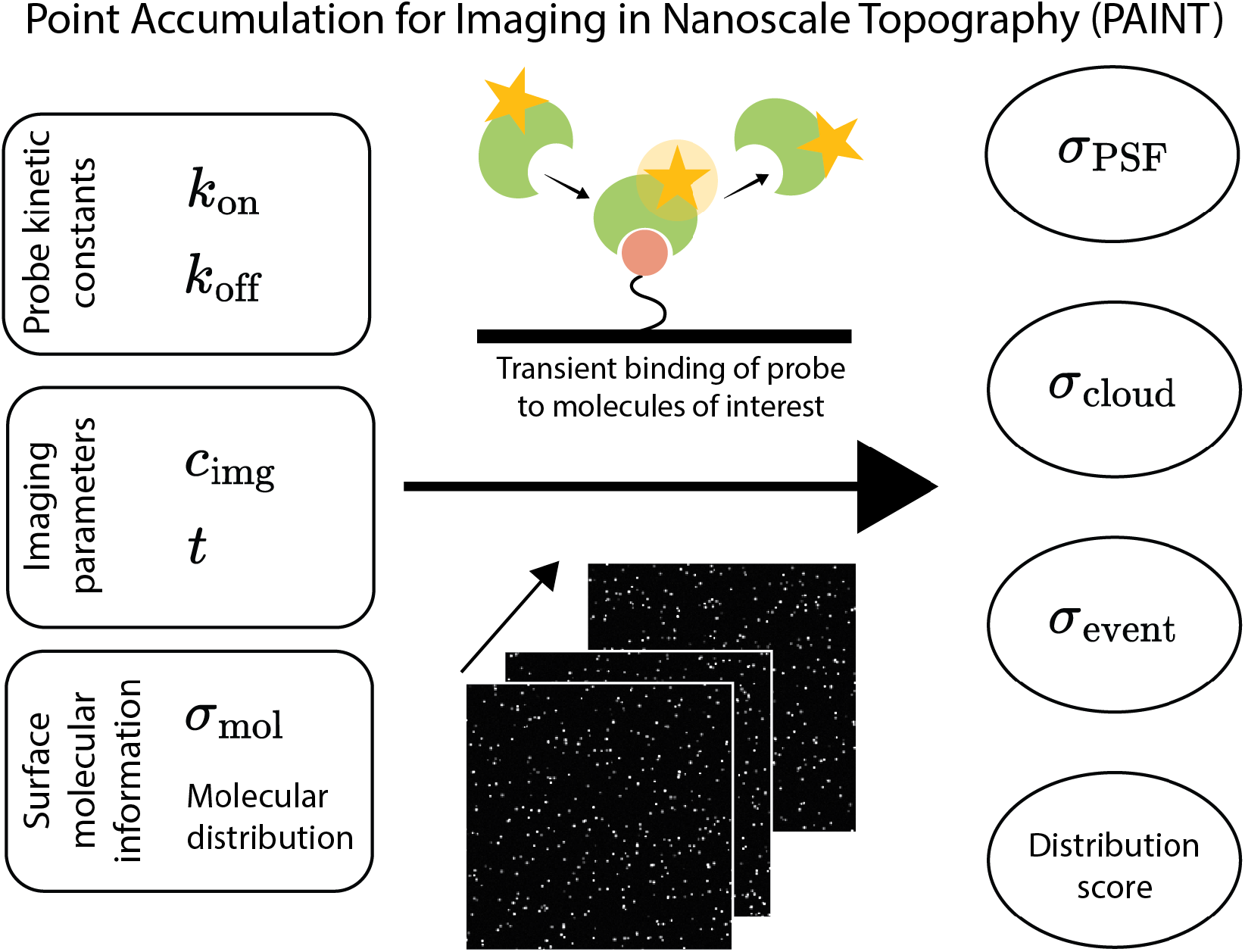
PAINT is a SMLM technique that achieves sub-diffraction imaging by temporally separating fluorescent emissions via the transient binding of dye-labeled probes to target molecules. The experimental or input parameters, namely the probe kinetic constants, imaging parameters, and true surface molecular information, collectively influence the PAINT output. Key output metrics obtained from the analysis of raw PAINT data include the point spread function (PSF) density *σ*_PSF_, localization cloud density *σ*_cloud_, binding event density *σ*_event_, and the observed molecular distribution score.

We define PSF detection accuracy using the F_1_ score, which accounts for the number of true positives (TP), false positives (FP), and false negatives (FN):

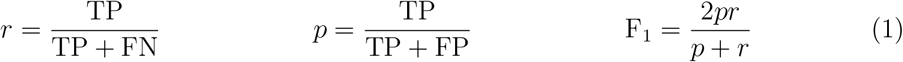

As shown in Equation 1, the F1 score reflects both recall *r*, the fraction of correctly identified PSFs, and precision *p*, the proportion of correct identifications among all localized PSFs. By varying the average PSF density 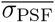 (averaged over all frames), we determined that PSFs can be detected with at least 90% accuracy up to a maximum density of 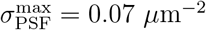, as shown in Figure 2(A).

**Figure 2.**
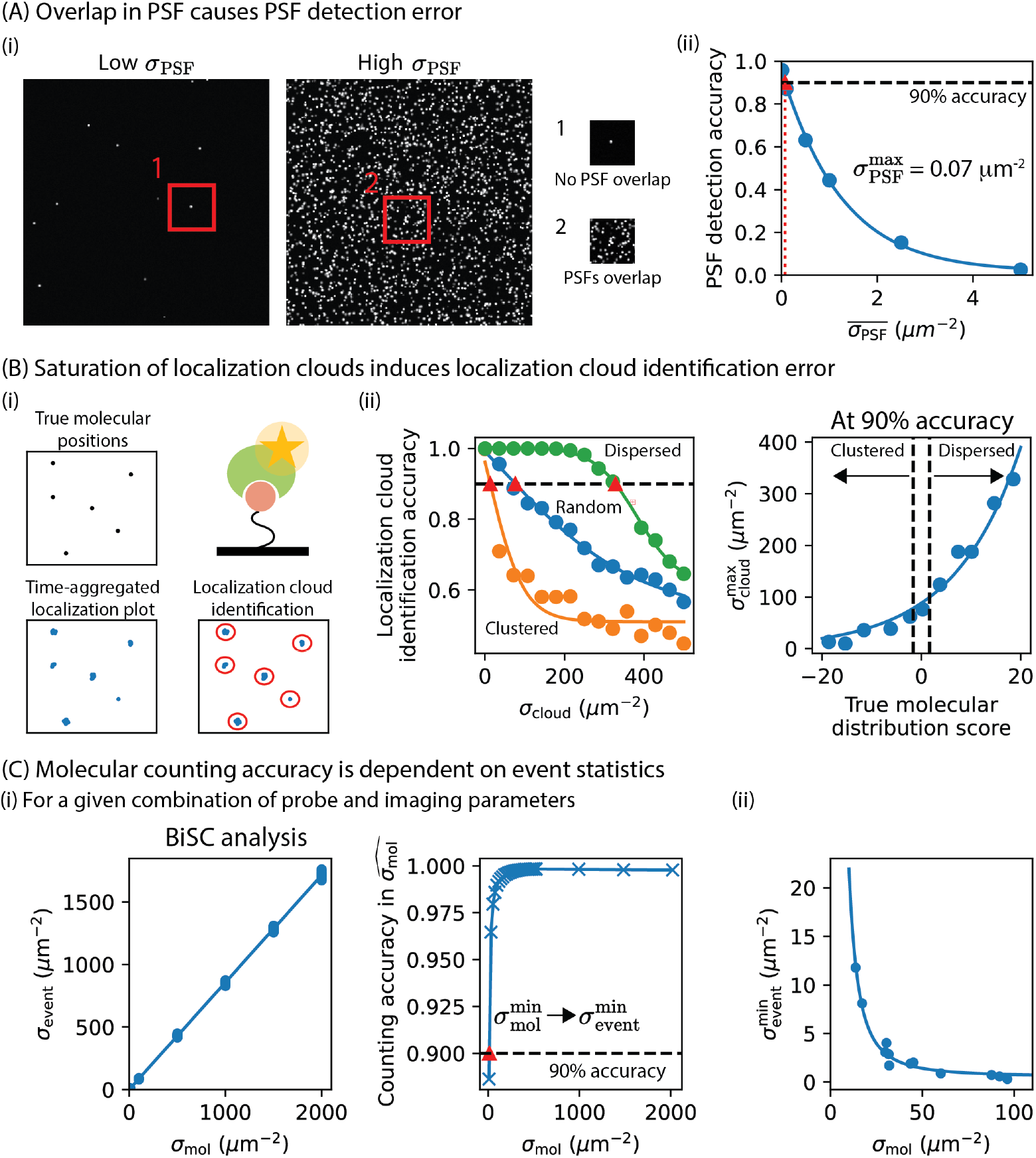
Molecular density quantification limits. (A) PSF saturation must be avoided to ensure accurate detection. An upper PSF density threshold of 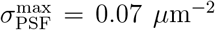 (red triangle) was identified, above which detection accuracy drops below 90%. (B) Localization cloud identification depends on both *σ*_cloud_ and molecular distribution. (i) Localization clouds were simulated from true molecular positions with defined spatial distributions. (ii) For each distribution score, the data are fitted with 1*/*(*a* + *e*^*bx*−*c*^) + *d* (solid lines), where *a, b, c* and *d* are fitting constants. 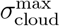 (red triangle) was determined at the 90% accuracy threshold. The relation between 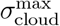 and distribution score is fitted with *ae*^*bx*^, where *a* and *b* are fitting constants. (C) BiSC-based quantification accuracy increases with event statistics. (i) For a given combination of probe and imaging parameters, a bootstrapping method was used to define 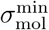 and 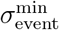 at 90% accuracy (red triangle). (ii) The relation between *σ*_mol_ and 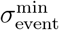, obtained across varying combination of probe and imaging parameters, is fitted with *a/x*^2^ + *b*, where *a* and *b* are fitting constants.

In SMLM, stacking localized PSFs across time results in a spatial map known as a time-aggregated localization plot. Because the bright time, the duration a PSF is visible, typically ranges from hundreds of microseconds to several seconds, a single molecule appears as a cloud of localizations. The underlying molecular position can be estimated from the centers of these localization clouds, which are identified using clustering algorithms such as meanshift clustering^17–19^or DBSCAN.^20,21^This step is critical both for direct molecular counting and for analyzing spatial distributions.

However, overlap between localization clouds, due to high cloud density *σ*_cloud_ or a clustered molecular distribution, can introduce errors in identifying individual localization clouds. To study this, we simulated ground truth molecular positions with predefined spatial distributions. A standardized distribution score, following the approach in previous work, ^14^was used to quantify spatial distributions, see Supplementary Section S1.2 for more details. Synthetic localizations were randomly generated around the true molecular positions and labeled accordingly, forming the ground-truth reference. We then applied mean-shift clustering to identify predicted localization clouds, assigning labels to each localization. The accuracy of this identification was quantified using the adjusted Rand index, which measures the similarity between ground-truth and predicted labels.^22^A perfect identification yields a score of 1.0, so we interpret the adjusted Rand index as the localization cloud identification accuracy.

As shown in Figure 2(B), identification accuracy decreases with increasing *σ*_cloud_, and the rate of decline depends strongly on the underlying molecular distribution. Specifically, for clustered distributions, accurate identification is only possible at relatively low *σ*_cloud_. We define the maximum identifiable cloud density 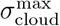 as the cloud density at which localization clouds can still be identified with at least 90% accuracy. We observed that 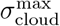 scales exponentially with the distribution score. In practice, this identification limit also depends on the localization precision and the bandwidth parameter used in clustering. In this study, we assume a typical localization precision of 8 nm and use an empirically optimized bandwidth parameter of 30 nm.^14^The effect of localization precision and bandwidth parameter on the 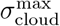 for varying spatial distributions is explored in Supplementary Section S2.

Furthermore, in high-density biofunctionalized surfaces, not all molecules are guaranteed to interact with dye-labeled probes within the finite measurement duration, leading to undersampling and undercounting when using direct counting. To address this, we previously developed the Biomolecule Sampling Compensation (BiSC) analysis,^14^a simulation-based method to compensate for sampling effects. In brief, we simulated binding events between probes and target molecules to explore the relationship between molecular density *σ*_mol_ and binding event density *σ*_event_ for various probe kinetic and imaging parameters. From the resulting linear relation, the experimentally observed *σ*_event_ can be used to estimate the molecular density 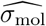. Supplementary Section S1.3 provides more details on the simulation procedure.

To assess the accuracy of this estimate, we applied a bootstrapping procedure: simulated datasets were resampled with replacement, and linear regression was repeated for each resampled set. Each regression model was used to estimate 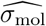 for a given *σ*_event_, repeated 1000 times. The relative residuals were calculated as follows:

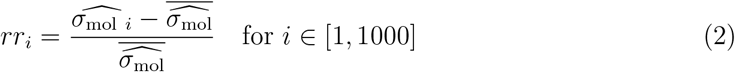

where *rr*_*i*_ denotes the relative residual and 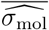 is the mean of the estimated densities. The counting accuracy is defined as the standard deviation of the relative residuals.

As shown in Figure 2(C), the counting accuracy improves with increasing *σ*_mol_, due to improved event statistics. Accordingly, we define the lower limit of quantifiable molecular density 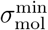 and the corresponding 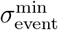 as the threshold below which 90% accuracy cannot be achieved. By repeating this analysis across multiple combinations of probe kinetic constants and imaging conditions (see Supplementary Table S4), we identified a scaling relationship between 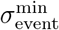 and (*σ*_mol_)^−2^. Thus, we interpret 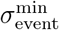 as the minimum event density required to achieve molecular quantification of a given *σ*_mol_ with at least 90% accuracy.

As previously discussed, the spatial distribution of biomolecules can be described and quantified using nearest-neighbor distance-based statistics. The reliability of this quantification depends on several factors: the precision of the estimated molecular positions, the molecular sampling ratio *s*_r_ (i.e., the fraction of molecules that interacted with the probe during the measurement), the true molecular density *σ*_mol_, and the underlying true spatial distribution. Here, we investigate how the observed distribution score varies with these factors. The data were generated as described in Figure 2(B)(i), where only a random subset of molecules was sampled according to sampling ratio *s*_r_:

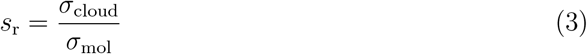

and localization clouds were generated around the sampled molecules.

Distribution scores were computed both for the true molecular positions and for the estimated positions,^14,23^using Equations 4 and 5, see Supplementary Section S1.2 for more details. Briefly, subsets of nearest neighbor distances (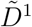, for 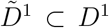) of size *m* were selected to ensure independence of the nearest neighbor distances. For each subset, the mean nearest neighbor distance 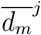 was calculated and standardized against the expected mean 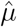 and standard error 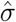 under a complete spatial randomness (CSR) hypothesis:

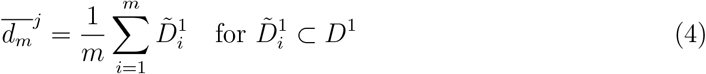

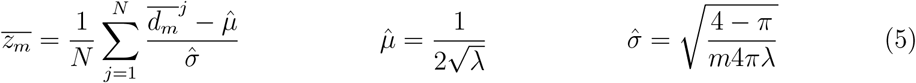

where *λ* denotes the molecular density. The standardized mean nearest-neighbor distance was calculated *N* times and subsequently averaged to obtain the distribution score. The resulting distribution score 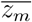 is compared against the CSR point pattern at the 5% significance level, *z*_0.05_ = 1.65, as illustrated in Figure 3(A).

**Figure 3.**
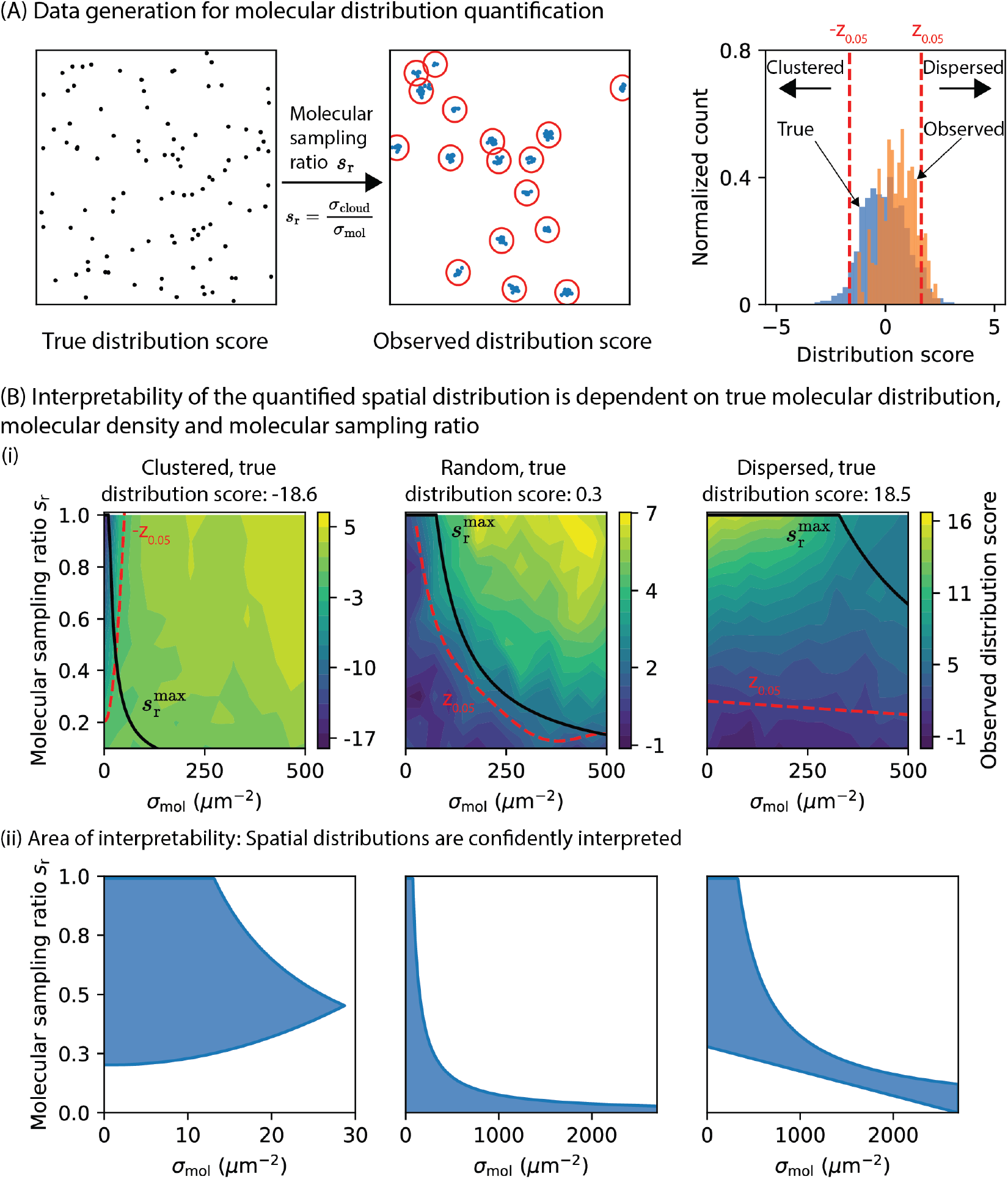
Interpretability of spatial molecular distribution quantification. (A) Molecular positions were simulated, followed by random sampling and generation of localization clouds around the sampled molecules. Distribution scores were computed for both the true and estimated positions. (B) The interpretability of the observed distribution score depends on the true spatial distribution, molecular density *σ*_mol_, and sampling ratio *s*_r_. (i) Contour maps of the observed score for three representative spatial distributions. (ii) Areas of interpretability showing combinations of *σ*_mol_ and *s*_r_ where the observed distribution score can be confidently interpreted.

Figure 3(B) shows three representative molecular distributions: clustered (true score = −18.6), random (true score = 0.3), and dispersed (true score = 18.5). For each case, the observed distribution score is plotted as a contour map over varying values of *σ*_mol_ and *s*_r_. For clustered and randomly distributed molecules, observed scores shift toward higher values (i.e., toward dispersion) as either *σ*_mol_ or *s*_r_ increases. For dispersed molecules, scores shift toward randomness as *s*_r_ decreases. Interpreting these contour maps is not straightforward. In practice, the true distribution is unknown, so the observed distribution score, calculated from the estimated molecular positions, is not an absolute value but an indicative one. Its meaning must be interpreted in reference to CSR, rather than to the ground truth. This raises an important question: how confidently can we interpret an observed distribution score (e.g., as clustered, random, or dispersed), based on the quantified values of *σ*_mol_ and *s*_r_?

Two conditions must be met for the interpretation to be valid. First, localization clouds must be identified with high accuracy to ensure precise estimation of the observed molecular positions. Based on the analysis in Figure 2(B), we define a maximum sampling ratio 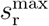 as

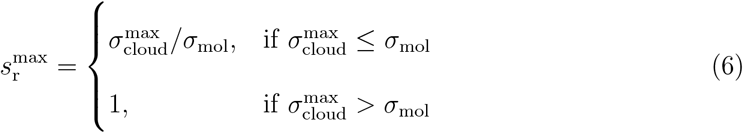

Note that 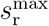 may exceed 1 in low-density regimes, indicating potential oversampling.

While the BiSC method is capable of handling both undersampling and oversampling, oversampling-related issues at low density are beyond the scope of this study. Therefore,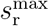 is capped at 1. Second, the observed distribution score must pass the statistical test against CSR. For example, if the underlying molecular distribution is clustered, the observed score must be less than −*z*_0.05_ (i.e., −1.65) to be interpreted as such.

These two conditions help clarify the structure of the contour maps in Figure 3(B). For randomly distributed molecules, the *z*_0.05_ contour line aligns with the boundary defined by 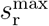, indicating that interpretability is governed primarily by the precision of estimated spatial positions of molecules. In contrast, for clustered and dispersed cases, both precision and statistical thresholding influence interpretability. Only a narrow region of *σ*_mol_ and *s*_r_ values allows reliable identification of clustering, whereas a much larger region supports reliable identification of dispersion. Figure 3(B)(ii) highlights the areas of interpretability, i.e., combinations of *σ*_mol_ and *s*_r_ for which the observed score can be reliably interpreted, for each of the three cases. For random and dispersed distributions, these regions are extrapolated to higher values of *σ*_mol_. In total, the analysis was performed across 11 true distribution scores ranging from −18 to 18. The corresponding evolution of the interpretability area is shown in the Supplementary Figure S3 and S4.

### 2.2 Global sensitivity analysis in PAINT output parameters

It is qualitatively known that the PAINT output, namely *σ*_PSF_, *σ*_cloud_, and *σ*_event_, is influenced by several factors, including the molecular association rate *k*_on_, dissociation rate *k*_off_, probe concentration *c*_img_, acquisition duration *t*, and the true molecular density *σ*_mol_. In this study, we systematically quantified the influence of these parameters using Monte Carlo simulations. Briefly, for a given *σ*_mol_, we simulated binding events between probe molecules and individual targets to generate binding time traces, as illustrated in Figure 4(A). This follows the same simulation framework as the BiSC analysis.^14^From these time traces, we computed the PAINT outputs using the following definitions:

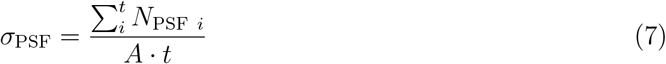

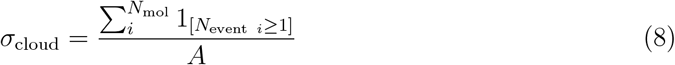

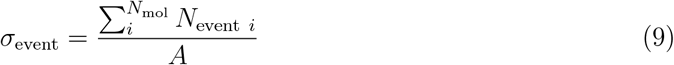

**Figure 4.**
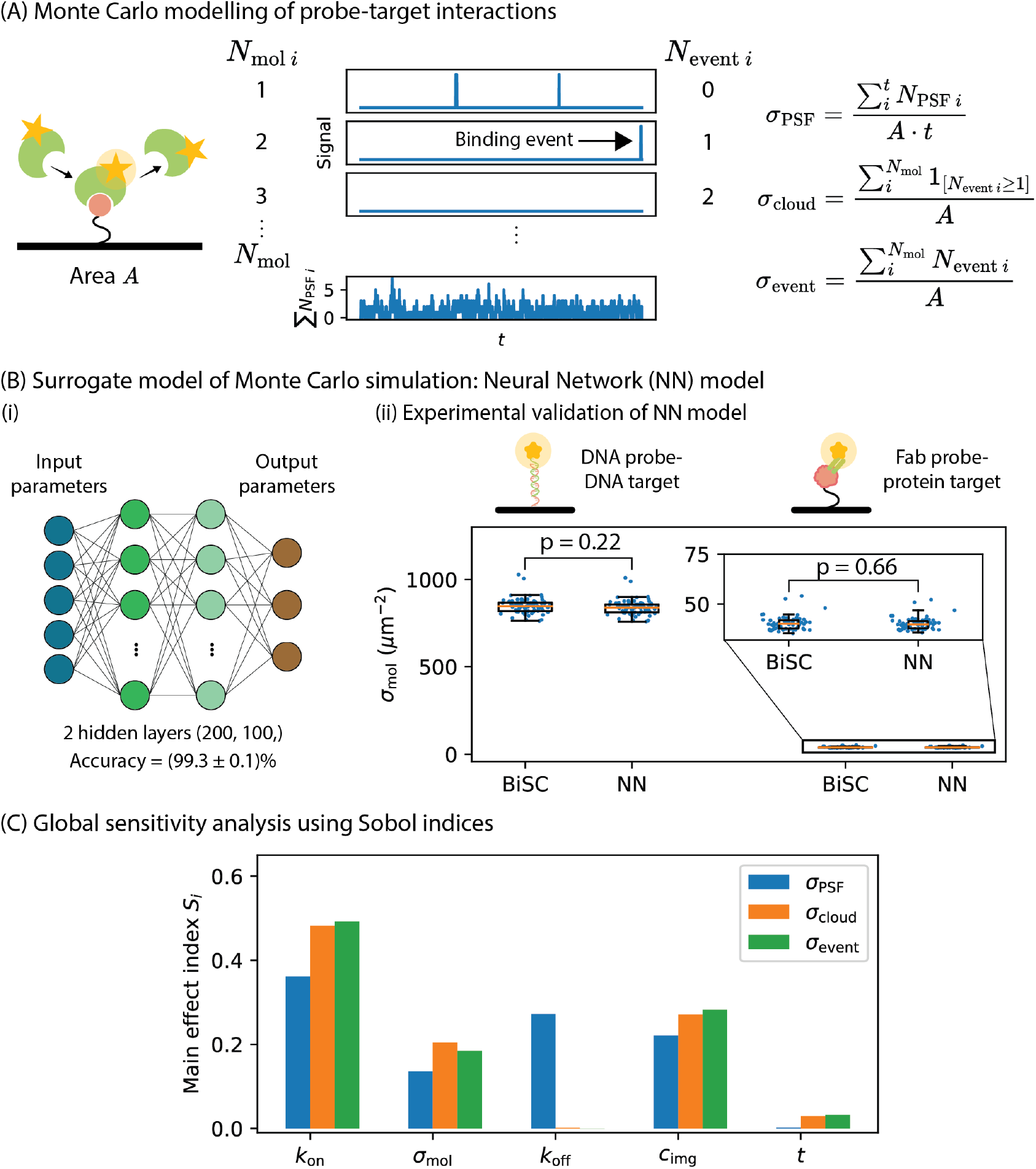
(A) Monte Carlo simulation of probe–target interactions in PAINT. Binding time traces are generated for each target molecule based on probe kinetics and imaging parameters. From these traces, three PAINT outputs are defined: *σ*_PSF_, *σ*_cloud_, and *σ*_event_. (B) Simulation data are used to train a neural network (NN) model. (i) The NN architecture includes two hidden layers with 200 and 100 nodes. The model achieves an accuracy of (99.3 *±* 0.1)%. (ii) Validation against BiSC-based analysis on experimental datasets shows no significant difference in estimated molecular densities (*p* > 0.05, Welch’s t-test). Molecular kinetic rates [*k*_on_ (M^−1^ s^−1^), *k*_off_ (s^−1^)] between the probe and target molecule are [10^6^, 1] and [3.6 × 10^6^, 1.9 × 10^−2^], for the ssDNA-based^14^and protein-based system,^24^respectively. (C) Main effect indices *S*_*i*_ are computed from global sensitivity analysis using the NN model.

Here, *σ*_PSF_ is defined as the mean number of PSFs observed over the acquisition duration *t* and area *A. σ*_cloud_ is the number of molecules with at least one binding event, normalized by *A*, and *σ*_event_ is the total number of binding events per area. Note that the spatial distribution of molecules was not explicitly modeled, as the interpretability of distribution quantification can already be inferred from the molecular sampling ratio *s*_r_ and *σ*_mol_.

To assess the influence of input parameters on PAINT output, we performed a global sensitivity analysis using Sobol indices.^25–27^These variance-based indices quantify the contribution of each input parameter to the total output variability. The first-order Sobol index *S*_*i*_, also denoted as the main effect index, captures the direct effect of a single parameter, while the total-order index *S*_*t*_ includes both the direct and all interaction effects. The difference *S*_*t*_ − *S*_*i*_ reflects the strength of parameter interactions. While Sobol indices offer a global perspective, estimating them reliably requires a large number of model evaluations, typically on the order of *N* (*k* + 2), where *N* is the number of samples per parameter and *k* is the number of parameters, resulting in thousands of simulations.

To reduce this computational cost, we trained a feedforward neural network (NN) as a surrogate model using data from the Monte Carlo simulations. ^28^The network architecture consists of one input layer, two hidden layers, and one output layer. We generated 3500 datasets by simulating probe-target interactions under different parameter combinations, see Supplementary Table S5. These were randomly split into 42% training, 28% validation, and 30% testing sets. The training data were used to optimize the model weights, while the validation data supported hyperparameter tuning and early stopping. Testing data provided an unbiased evaluation of model performance. The training of the model is shown in Supplementary Figure S1.

Since neural networks are inherently stochastic, due to randomness in initialization and optimization, we evaluated model consistency in two ways: (i) training the model 100 times using the same data and varying random seeds, and (ii) splitting the dataset 100 times and training with a fixed seed. Model accuracy was evaluated using the coefficient of determination *R*^2^ and the average accuracy was found to be (99.3 ± 0.1)%. To further validate the NN model, we used it to estimate molecular densities from two experimental datasets: one ssDNA-functionalized surface probed by ssDNA probes,^14,29,30^and one protein-functionalized surface probed by Fab fragments.^24^Results were compared to BiSC-based estimates, and no significant difference was found based on Welch’s t-test, confirming that the NN model reproduces the Monte Carlo results (Figure 4(B)).

Figure 4(C) presents the main effect indices *S*_*i*_ of each input parameter on the PAINT outputs, as derived from sensitivity analysis on the NN model. Total-order indices *S*_*t*_ are provided in the Supplementary Table S8. In general, the PAINT outputs were strongly dominated by main effects, indicating limited interaction between parameters. Among all inputs, the molecular association rate *k*_on_ had the largest influence, followed by probe concentration *c*_img_ and molecular density *σ*_mol_, across all outputs. For *σ*_PSF_, however, *k*_off_ contributed more than *σ*_mol_, which is expected since longer binding durations (lower *k*_off_) increase the probability of PSF detection per measurement frame. Interestingly, acquisition duration *t* had the smallest effect (after *k*_off_) on both *σ*_cloud_ and *σ*_event_, even though longer acquisition times are intuitively expected to increase the number of detectable binding events. While the qualitative effects of experimental parameters on PAINT output are generally understood, this sensitivity analysis quantitatively ranks their relative importance and highlighted the dominant role of probe kinetics and concentration.

### 2.3 Rational design of PAINT experiment and probe

In a PAINT experiment, the goal is to quantify both the spatial density and distribution of surface-bound molecules. However, as shown in previous sections, these two properties also influence the accuracy and interpretability of the quantification itself. Ideally, one would select probe and imaging parameters that maximize the likelihood of obtaining accurate and interpretable spatial molecular information. To investigate this, we used an in silico approach to generate PAINT outputs over a wide range of input parameters using the trained neural network (NN) model (see Table 1). Note that the true distribution score was not explicitly simulated, as its influence is implicitly captured through the combination of *σ*_mol_ and the molecular sampling ratio *s*_r_. The quantification limits derived in Figure 2 and 3 were then applied to assess the viability of molecular quantification under each condition. The effect of each quantification limit on the viability of molecular quantification is discussed in Supplementary Section S4.3.

**Table 1:** The input parameters used to generate PAINT outputs via the NN model, sampled uniformly within defined ranges [min,max] for N values per parameter

|  | Min | Max | # samples |
| --- | --- | --- | --- |
| $k_{\text{on}}$ ($\text{M}^{-1} \text{s}^{-1}$) | $10^3$ | $10^7$ | 25 |
| $k_{\text{off}}$ ($\text{s}^{-1}$) | $10^{-4}$ | $10^0$ | 25 |
| $c_{\text{img}}$ (M) | $10^{-12}$ | $10^{-8}$ | 100 |
| $t$ (min) | 10 | 10 | 50 |
| $\sigma_{\text{mol}}$ ($\mu\text{m}^{-2}$) | 0.1 | 2700 | 300 |
| True distribution score | -18 | 18 | 11 |

We first considered the effect of the probe concentration *c*_img_ on quantification performance for a DNA probe with reference kinetics *k*_on_ = 10^6^ M^−1^ s^−1^ and *k*_off_ = 1 s^−1^ (Figure 5(A)). Because acquisition duration *t* has limited influence on the PAINT output, we focused on the effect of *c*_img_. Two scenarios were analyzed: (i) quantifying molecular density only, and (ii) quantifying both density and spatial distribution. For each value of *c*_img_ and for each true molecular distribution, we computed the range of *σ*_mol_ values that could be quantified with at least 90% accuracy. This range was normalized by the full simulated *σ*_mol_ range (*≈* 2700 *µ*m^−2^), yielding a maximum normalized value of 1, see Supplementary Section S4.1 for more details. This gives a map of quantifiable density range for a given probe.

**Figure 5.**
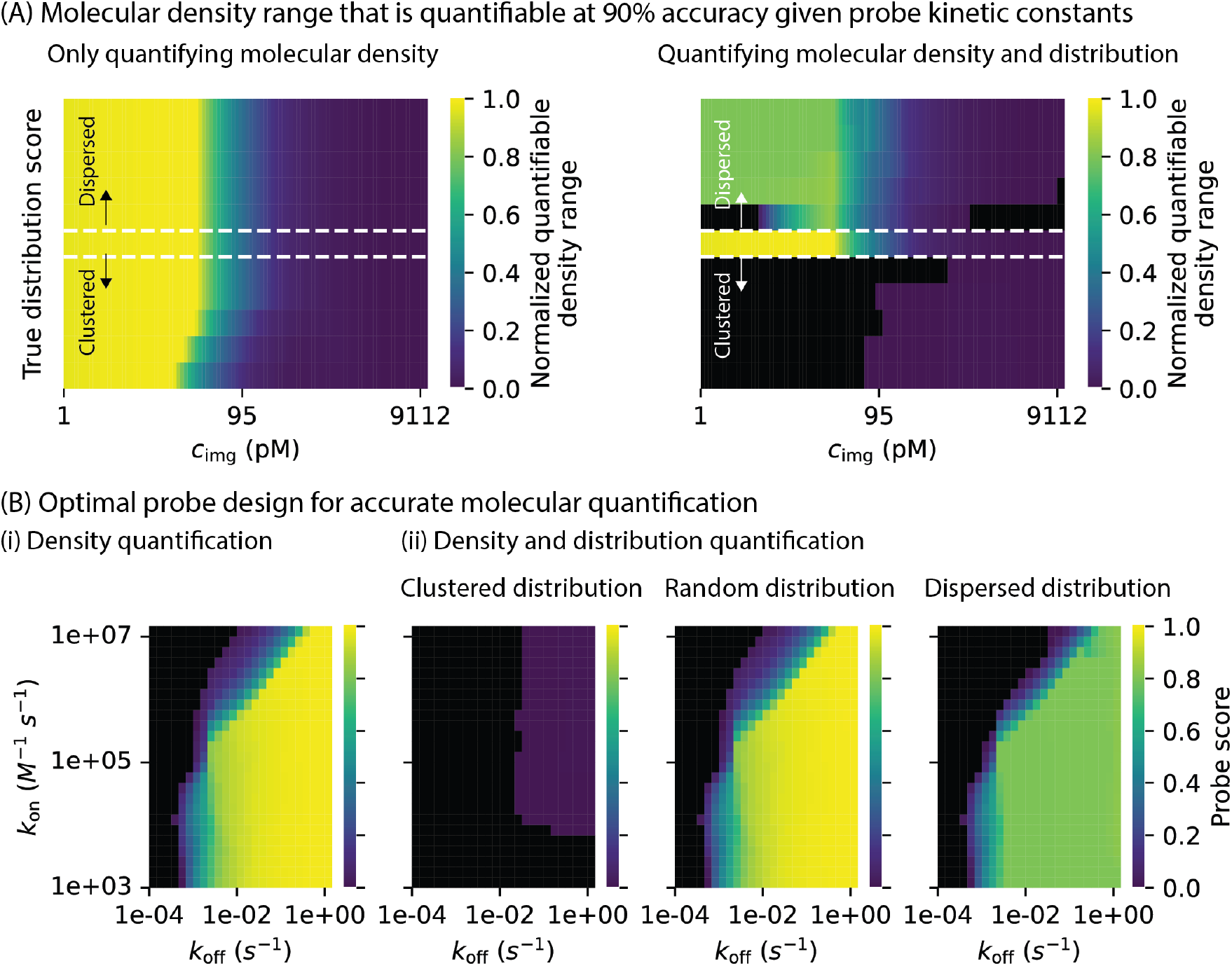
(A) For a given set of probe kinetic constants, contour maps show the range of molecular densities that can be quantified with at least 90% accuracy. Two scenarios are evaluated: density-only quantification and joint quantification of density and spatial distribution. The white dotted lines serve as a visual guide; deviations from it reflect increasing molecular clustering or dispersity. Regions in between the white dotted lines indicate randomly distribution molecules. Black regions indicate parameter combinations where no density can be reliably quantified. (B) Optimal probe design maps for (i) precise density quantification and (ii) combined density and distribution quantification. Black regions denote probes that fail to support high-accuracy quantification under the tested conditions.

When considering only density quantification, lowquantified with high accuracy values of *c*_img_ enable precise quantification across the entire simulated *σ*_mol_ range. However, when spatial distribution quantification is included as an additional criterion, only the trend for randomly distributed molecules resembles that of the density-only case. This is because the interpretability of distribution scores primarily depends on the accuracy with which individual localization clouds are identified. For dispersed distributions, a broad range of *σ*_mol_ can still be quantified at low *c*_img_ values, owing to the wide interpretability regions. However, the overall quantifiable range is slightly reduced, as low-density systems do not meet the minimum molecular sampling ratio required for statistically interpreting the distribution as dispersed. This threshold is governed by the statistical criterion that the distribution score must exceed *z*_0.05_. In contrast, for clustered distributions, only high values of *c*_img_ allowed even limited quantification, reflecting the inherently narrow area of interpretability for clustered spatial arrangements. Black squares in Figure 5(A) indicate parameter combinations where no *σ*_mol_ values could be reliably quantified.

In practice, researchers are often interested not only in molecular density, but also in spatial distribution. The findings here highlight a critical limitation: spatial distributions that are inherently clustered cannot be effectively quantified using PAINT under high-density conditions. This raises an important consideration: what is the actual molecular feature of interest? For example, in a high-density surface functionalized via a multivalent linker such as neutravidin, clustering arises naturally due to multiple binding sites. In such cases, studying the spatial distribution of the linker (e.g., neutravidin) rather than the conjugated molecule may be more relevant. Since linkers are typically of lower density, their spatial distributions are more likely to fall within interpretable limits. Thus, while PAINT is limited in resolving high-density, clustered distributions, this limitation may not be critical if the system’s design and biological context are appropriately considered. In cases where it is essential to quantify the spatial distribution of high-density, clustered molecules, the interpretability region must be expanded. This can be achieved by improving resolution (e.g., via RESI), and ensuring high molecular sampling ratios (see Supplementary Figure S5).

In the previous section, we emphasized the importance of probe kinetic constants on the PAINT output. Figure 5(B) shows the parameter space over which different probes can yield high-accuracy quantification, along with the corresponding *σ*_mol_ ranges they can reliably probe. For molecular density quantification, we computed the mean normalized quantifiable *σ*_mol_ range (averaged over the number of true distributions considered) across the *c*_img_ values for each probe, and defined the probe score as the maximum of the means. When both density and spatial distribution quantification are required, the map is divided into three spatial regimes, namely clustered, random, and dispersed. For each regime, the mean normalized quantifiable density range is calculated across *c*_img_, and the maximum value is taken as the probe score. The probe score thus represents the fraction of the full *σ*_mol_ range that can be quantified with at least 90% accuracy within the simulated *c*_img_ range, and reflects the objective of identifying probes that support reliable quantification over a broad range of true molecular densities *σ*_mol_ and spatial distributions, regardless of the probe concentration used. Supplementary Figure S7 illustrates the calculation of probe scores reported in this work.

When considering only density quantification, a wide range of probe kinetic constants, specifically, *k*_on_ values from 10^3^ to 10^7^ M^−1^ s^−1^ and *k*_off_ values from 10^−2^ to 1 s^−1^, support accurate quantification across broad *σ*_mol_ ranges. This flexibility is encouraging for the development of novel PAINT-compatible probes, as it suggests that a wide range of kinetic parameters can still yield reliable density estimates. When both density and spatial distribution quantification are considered, a similarly broad range of probe kinetics supports accurate and interpretable quantification for randomly and dispersed distributed systems. However, for dispersed systems, the quantifiable *σ*_mol_ range is slightly reduced because molecular sampling ratio thresholds required for statistical interpretability are not met for low-density systems. An important observation is that probes with similar probe scores may operate in different *c*_img_ regimes, depending on their kinetic parameters. This variability is illustrated in Supplementary Figure S9 and S10, and emphasizes the need to tailor imaging conditions to specific probe kinetics.

In contrast, for clustered distributions, only a narrow range of probe kinetics enables limited quantification, again highlighting the intrinsic challenge of interpreting high-density clustered spatial arrangements. Improving PAINT resolution slightly expands the usable kinetic space and increases the quantifiable *σ*_mol_ range, as shown in Supplementary Figure S11. However, the range remains narrower than that for random or dispersed systems, indicating a need for either further enhancement in PAINT resolution or new analytical strategies to account for limitations in distribution quantification. One promising direction is machine-learning-based analysis that learns the relationship between PAINT precision, true molecular densities, sampling ratio, and distribution scores. Such a model could enable inverse estimation of true molecular distributions, even under suboptimal resolution. Overall, these findings provide insight into the range of probe kinetics compatible with highaccuracy molecular quantification in PAINT, offering practical guidance for the design of next-generation, non-DNA-based PAINT probes.

## Conclusions

We presented a systematic, simulation-guided study to investigate how probe kinetic constants, imaging parameters, and surface molecular characteristics influence PAINT output metrics for accurate molecular quantification. By defining quantification limits for key PAINT outputs, namely PSF density *σ*_PSF_, localization cloud density *σ*_cloud_, and binding event density *σ*_event_, we established the conditions under which molecular density can be quantified with high accuracy (≥ 90%), and identified when spatial distribution quantification can be reliably interpreted. Through a neural network (NN) model trained on Monte Carlo simulation data, we performed a global sensitivity analysis using Sobol indices to quantify the contribution of each input parameter to the variability of PAINT outputs. This revealed the dominant roles of probe association rate *k*_on_, dissociation rate *k*_off_, and probe concentration *c*_img_, while highlighting minimal contributions from acquisition duration *t* under the simulated conditions.

Leveraging the trained NN model as a surrogate, we explored the parameter space across a wide range of experimental conditions to identify optimal regimes that enable accurate and interpretable molecular quantification. Our results underscore that while a broad range of kinetic and concentration parameters can support high-accuracy density quantification, the reliable interpretation of spatial distributions is more restrictive, particularly for highdensity and clustered molecular systems. We also emphasize that interpretability depends also on a priori knowledge of the molecular system of study, which can help redirect the focus of analysis toward more tractable molecular targets, such as linker molecules. When highdensity clustered distributions must be characterized, improvements in PAINT resolution or advanced analytical corrections become essential.

Overall, the described in silico approach provides a quantitative foundation for selecting experimental conditions that ensure accurate molecular quantification in PAINT experiments. Beyond optimizing imaging parameters, this framework offers practical design guidelines for developing next-generation PAINT-compatible imaging probes, especially those targeting non-DNA systems. By clarifying the interplay between experimental variables and quantification limits, this work contributes to expanding the applicability and reliability of PAINT for molecular quantification across diverse biological and synthetic surfaces.

## Data availability

The data supporting this article have been included as part of the Supplementary Information.

## Acknowledgements

This work has received funding from the European Union’s Horizon 2020 research and innovation program under the Marie Skłodowska-Curie grant agreement No. 955623 (H2020-MSCA ITN-CONSENSE). Part of this work was funded by The Netherlands National Growth Fund Programme NXTGEN HighTech. We thank Claire Michielsen and Junhong Yan for providing materials essential for the PAINT experiments on protein-functionalized surfaces.

