## Supplementary Information for "Simulation-Guided Exploration of PAINT Parameter Space for Accurate Molecular Quantification"

### Contents

**S1 Simulation procedure** **S-3**

S1.1 Point spread function (PSF) density . . . . . . . . . . . . . . . . . . . . . . . . S-3

S1.2 Localization cloud density and spatial distribution . . . . . . . . . . . . . . . . S-4

S1.3 Binding event density . . . . . . . . . . . . . . . . . . . . . . . . . . . . . . . . S-5

S1.4 Neural network surrogate model . . . . . . . . . . . . . . . . . . . . . . . . . . S-8

**S2 Factors affecting localization cloud quantification limit** **S-9**

**S3 Areas of interpretability in quantified distribution scores** **S-10**

**S4 Probe scores** **S-15**

S4.1 Range of molecular density quantifiable with at least 90% accuracy per probe S-15

S4.2 Calculation of probe scores . . . . . . . . . . . . . . . . . . . . . . . . . . . . . S-16

S4.3 Effect of quantification limit and probe kinetics . . . . . . . . . . . . . . . . . S-18

S4.4 Effect of PAINT resolution . . . . . . . . . . . . . . . . . . . . . . . . . . . . . S-23

**S5 Additional information** **S-25**

S5.1 Experimental details for NN experimental validation dataset . . . . . . . . . S-25

S5.1.1 Materials and chemicals . . . . . . . . . . . . . . . . . . . . . . . . . . S-25

S5.1.2 Coverslip surface functionalization . . . . . . . . . . . . . . . . . . . . S-26

S5.1.3 PAINT imaging and analysis . . . . . . . . . . . . . . . . . . . . . . . . S-26

S5.2 Sobol analysis results . . . . . . . . . . . . . . . . . . . . . . . . . . . . . . . . S-28

**References** **S-29**

### S1 Simulation procedure

#### S1.1 Point spread function (PSF) density

To investigate the effect of point spread function (PSF) density $\sigma_{\text{PSF}}$ on PSF detection accuracy, the ThunderSTORM simulation engine was used to generate PSFs over 100 frames for a given density.[S1] The input parameters used in the simulations are listed in Table S1. The generated PSFs serve as ground truth positions. Simulated raw data were analyzed using ThunderSTORM to localize PSFs in each frame. The resulting localizations were then compared to the ground truth and evaluated in terms of true positives (TP), false positives (FP), and false negatives (FN), using the following metrics:

$$r = \frac{\text{TP}}{\text{TP} + \text{FN}} \qquad p = \frac{\text{TP}}{\text{TP} + \text{FP}} \qquad \text{F}_1 = \frac{2pr}{p + r} \tag{S1}$$

Here, recall $r$ quantifies the fraction of ground truth PSFs correctly detected, while precision $p$ reflects the proportion of correct detections among all localized PSFs. The $\text{F}_1$ score combines both into a single performance metric, where values near 0 indicate poor detection performance and values close to 1 indicate high accuracy. In this study, the $\text{F}_1$ score is used as a measure of PSF detection accuracy.

Table S1: Input values for ThunderSTORM simulation engine.

| | | |
|---|---|---|
| Field of view width | ($\mu$m) | 30 |
| Field of view height | ($\mu$m) | 30 |
| Frames | | 100 |
| PSF width | (nm) | 100 to 200 |
| PSF intensity | (photon) | 400 to 20000 |
| Mean photon background | (photon) | 30 |
| PSF density $\sigma_{\text{PSF}}$ | ($\mu\text{m}^{-2}$) | 0.01 to 2.5 |

#### S1.2 Localization cloud density and spatial distribution

To investigate the effect of localization cloud density $\sigma_{\text{cloud}}$ on cloud identification accuracy and to examine the factors influencing quantified spatial distribution, spatial point patterns were simulated with a given molecular density $\sigma_{\text{mol}}$ and a predefined spatial distribution, characterized by an average nearest-neighbor distance $\overline{D^1}$. A subset of molecular positions was randomly selected according to a specified molecular sampling ratio $s_{\text{r}}$, and localizations were generated around these sampled positions using a Gaussian distribution with a standard deviation of 8 nm, reflecting experimentally measured localization uncertainty.[S2] The number of localizations generated are sampled from an exponential distribution with a characteristic bound state lifetime of 1 s, corresponding to a molecular dissociation rate $k_{\text{off}}$ of 1 s$^{-1}$. For evaluating localization cloud identification accuracy, full sampling ($s_{\text{r}} = 1$) was assumed. Ground-truth labels were assigned to each simulated localization. Mean-shift clustering with a bandwidth of 30 nm was applied to identify localization clouds. The adjusted Rand index was used to compare the clustering results with ground-truth labels, and is referred to as the localization cloud identification accuracy.[S3] The input values for generating the simulated data is summarized in Table S2.

To assess spatial distribution for both true and estimated molecular positions, a nearest-neighbor distance-based statistical test—the Clark–Evans test—was applied to evaluate deviations from complete spatial randomness (CSR).[S4] The test compares the observed average nearest-neighbor distance $\overline{D^1}$ to the expected value $\hat{\mu}$ under CSR and determines whether the pattern is significantly clustered or dispersed. To mitigate edge effects, nearest-neighbor distances from points within the outer 10% boundary of the region of interest were excluded from the statistics. To satisfy the independence assumption of $D^1$, $N$ subsets of size $m$ were randomly sampled from the nearest-neighbor distance set to compute mean values:

$$\overline{d_m}^j = \frac{1}{m}\sum_{i=1}^{m} \tilde{D}_i^1 \quad \text{for } \tilde{D}_i^1 \subset D^1 \tag{S2}$$

With this, the standardized mean nearest neighbor distance $\overline{z_m}$, also denoted as the distribution score, can be computed via

$$\overline{z_m} = \frac{1}{N}\sum_{j=1}^{N}\frac{\overline{d_m}^{j} - \hat{\mu}}{\hat{\sigma}} \qquad \hat{\mu} = \frac{1}{2\sqrt{\lambda}} \qquad \hat{\sigma} = \sqrt{\frac{4-\pi}{m4\pi\lambda}} \tag{S3}$$

where $\lambda$ denotes the point density, $\hat{\mu}$ the expected mean nearest-neighbor distance, $\hat{\sigma}$ the standard deviation of the expected mean nearest-neighbor distance. The distribution score is tested against the distribution score of a CSR point pattern at a 5% significance level $z_{0.05} = 1.65$, giving

$$\begin{aligned} \overline{z_m} &< -z_{0.05} &&: \text{Significant clustering} \\ \overline{z_m} &> \quad z_{0.05} &&: \text{Significant dispersion} \end{aligned} \tag{S4}$$

Table S2: Input values to generate simulated data. *Note that, when studying localization cloud identification accuracy, $s_{\mathrm{r}}$ is set to 1.

| | | Values |
|---|---|---|
| True molecular density $\sigma_{\mathrm{mol}}$ | ($\mu$m$^{-2}$) | 0.1 to 500 |
| True distribution score | | -18 to 18 |
| Molecular sampling ratio $s_{\mathrm{r}}$ | | 0.1 to 1* |
| Average localization uncertainty | (nm) | 8 |
| Analysis search region | (nm) | 30 |
| $k_{\mathrm{off}}$ | (s$^{-1}$) | 1 |

#### S1.3 Binding event density

In previous work, a simulation-based method known as Biomolecule Sampling Compensation (BiSC) analysis was developed to estimate surface molecular density from experimentally observed binding event density.[S2] BiSC compensates for the effect of biomolecule undersampling in molecular quantification by simulating binding dynamics using Monte Carlo methods.

In this approach, signal time traces for each biomolecule within a region of interest of area $A$ and acquisition duration $t$ are simulated. Each time trace consists of alternating bound and unbound states, with lifetimes governed by the molecular association rate $k_{\text{on}}$, dissociation rate $k_{\text{off}}$, and probe concentration $c_{\text{img}}$. The characteristic lifetimes are defined as:

$$\tau_{\text{b}} = \frac{1}{k_{\text{off}}} \qquad \tau_{\text{ub}} = \frac{1}{k_{\text{on}} \cdot c_{\text{img}}} \tag{S5}$$

where $\tau_{\text{b}}$ and $\tau_{\text{ub}}$ denote the characteristic bound and unbound state lifetime respectively. Bound and unbound durations are sampled from exponential distributions with the corresponding lifetimes. The probability of a molecule being in the bound state is given by:

$$\text{P(bound)} = \frac{c_{\text{img}}}{c_{\text{binder}} + K_{\text{d}}} \qquad \text{with } K_{\text{d}} = \frac{k_{\text{off}}}{k_{\text{on}}} \tag{S6}$$

where $c_{\text{binder}}$ is the effective volumetric biomolecule concentration and $K_{\text{d}}$ is the equilibrium dissociation constant. The binding probability, P(bound), depends on both the probe concentration and hybridization kinetics (see Table S3). Time traces are generated with a 100 ms integration time. The total number of binding events observed per trace is summed and normalized by the area $A$ to compute the binding event density $\sigma_{\text{event}}$. By varying $\sigma_{\text{mol}}$, a linear relationship between $\sigma_{\text{mol}}$ and $\sigma_{\text{event}}$ is established. Each simulation is repeated 20 times per $\sigma_{\text{mol}}$ value, with parameter details summarized in Table S4.

Table S3: Dependence of the binding probability on imager strand concentration and hybridization kinetics.

| $c_{\text{binder}}$ | $K_{\text{d}}$ | P(bound) |
|---|---|---|
| $\ll c_{\text{img}}$ | $\ll c_{\text{img}}$ | 1 |
| $\ll c_{\text{img}}$ | $\gg c_{\text{img}}$ | $c_{\text{img}}/K_{\text{d}}$ |
| $\gg c_{\text{img}}$ | $\ll c_{\text{binder}}$ | $c_{\text{img}}/c_{\text{binder}}$ |
| $\gg c_{\text{img}}$ | $\gg c_{\text{binder}}$ | $c_{\text{img}}/K_{\text{d}}$ |

To assess the accuracy of the estimated molecular density $\widehat{\sigma_{\text{mol}}}$, a bootstrapping procedure is applied: simulated datasets are resampled with replacement, and linear regression is performed on each resampled set. Each model is then used to predict $\widehat{\sigma_{\text{mol}}}$ for a fixed $\sigma_{\text{event}}$,

Table S4: Input parameters for BiSC analysis.

| | | Value |
|---|---|---|
| $k_{\text{off}}$ | $(\text{s}^{-1})$ | 0.1, 1, 10 |
| $k_{\text{on}}$ | $(\text{M}^{-1}\ \text{s}^{-1})$ | $5 \times 10^4$ to $10^7$ |
| $c_{\text{img}}$ | (M) | $2.5 \times 10^{-11}$ |
| $t$ | (min) | 30, 60, 90 |
| Integration time | (s) | 0.1 |
| $\sigma_{\text{mol}}$ | $(\mu\text{m}^{-2})$ | 10 to 2000 |

repeated 1000 times. Relative residuals are computed as:

$$rr_i = \frac{\widehat{\sigma_{\text{mol}}}_i - \overline{\widehat{\sigma_{\text{mol}}}}}{\overline{\widehat{\sigma_{\text{mol}}}}} \quad \text{for } i \in [1, 1000] \tag{S7}$$

where $rr_i$ denotes the relative residual and $\overline{\widehat{\sigma_{\text{mol}}}}$ is the mean of the estimated densities. The standard deviation of the relative residuals is taken as the measure of counting accuracy. In Figure 2(C)(ii), varying combination of probe kinetic constants and imaging parameters are used to derive the relationship between $\sigma_{\text{mol}}$ and the minimum binding event density $\sigma_{\text{event}}^{\text{min}}$ required for molecular quantification accuracy of at least 90%. Table S5 summarizes the conditions used.

Table S5: Combinations of probe kinetic constants and acquisition duration used in Figure 2(C)(ii).

| $\sigma_{\text{mol}}$ $(\mu\text{m}^{-2})$ | $\sigma_{\text{event}}^{\text{min}}$ $(\mu\text{m}^{-2})$ | $k_{\text{on}}$ $(\text{M}^{-1}\ \text{s}^{-1})$ | $k_{\text{off}}$ $(\text{s}^{-1})$ | $t$ (min) |
|---|---|---|---|---|
| 15.23 | 13.17 | $10^7$ | 1 | 60 |
| 16.60 | 7.83 | $5 \times 10^6$ | 1 | 60 |
| 28.64 | 2.97 | $10^6$ | 0.1 | 60 |
| 30.42 | 4.00 | $10^6$ | 1 | 60 |
| 31.53 | 1.68 | $10^6$ | 1 | 90 |
| 32.33 | 2.92 | $10^6$ | 10 | 60 |
| 42.25 | 1.85 | $10^6$ | 1 | 30 |
| 45.61 | 2.03 | $5 \times 10^5$ | 1 | 60 |
| 59.83 | 0.89 | $2 \times 10^5$ | 1 | 60 |
| 86.31 | 0.71 | $10^5$ | 1 | 60 |
| 92.06 | 0.58 | $7 \times 10^4$ | 1 | 60 |
| 93.34 | 0.29 | $5 \times 10^4$ | 1 | 60 |

#### S1.4 Neural network surrogate model

To reduce computational cost, a neural network (NN) surrogate model was trained using data generated from the Monte Carlo simulations described in the previous section.[S5] The simulation outputs include point spread function density $\sigma_{\mathrm{PSF}}$, localization cloud density $\sigma_{\mathrm{cloud}}$, and binding event density $\sigma_{\mathrm{event}}$. A total of 3500 datasets, corresponding to different combinations of input parameters (see Table S6), were used to train the NN model. The neural network architecture consists of two hidden layers with 200 and 100 neurons, respectively. The rectified linear unit (ReLU) was used as the activation function in the hidden layers.[S6] No explicit regularization was applied during training.

Table S6: The input parameters used to generate PAINT outputs via the NN model were sampled uniformly within defined ranges [min,max] for N values per parameter.

| | | Min | Max | # samples |
|---|---|---|---|---|
| $k_{\mathrm{on}}$ | $(\mathrm{M}^{-1}\ \mathrm{s}^{-1})$ | $10^3$ | $10^7$ | 5 |
| $k_{\mathrm{off}}$ | $(\mathrm{s}^{-1})$ | $10^{-}4$ | $10^0$ | 5 |
| $c_{\mathrm{img}}$ | (M) | $10^{-12}$ | $10^{-9}$ | 4 |
| $t$ | (min) | 10 | 110 | 5 |
| $\sigma_{\mathrm{mol}}$ | $(\mu\mathrm{m}^{-2})$ | 10 | 2700 | 7 |

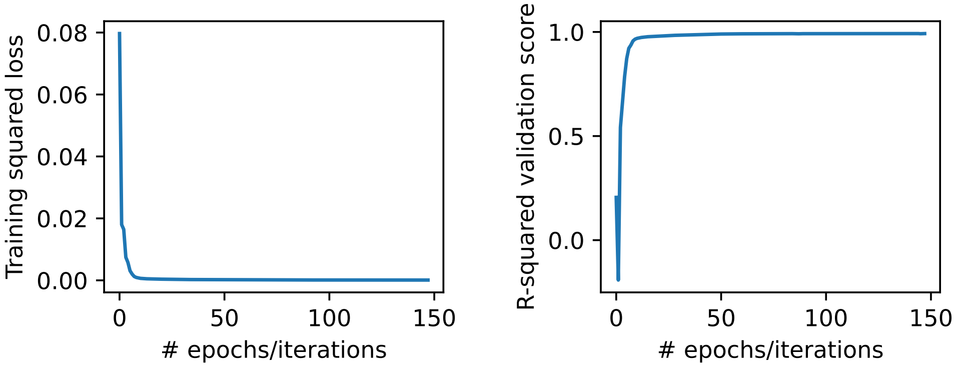


Figure S1: The training loss (left), defined as mean squared error, decreases consistently with increasing epochs, indicating effective model fitting. The validation score (right), measured by the coefficient of determination ($R^2$), remains high throughout and plateaus at later stages, confirming that the neural network was successfully trained without overfitting.

Figure S1 shows the training progression, where the loss, defined as the mean squared error, decreases over epochs. Validation performance was tracked using the coefficient of determination $R^2$, defined as:

$$R^2 = 1 - \frac{\sum (y_i - \hat{y}_i)^2}{\sum (y_i - \overline{y})^2} \tag{S8}$$

As training progresses, $R^2$ improves and eventually plateaus. In cases of overfitting, the validation score may decline during later stages of training.

### S2 Factors affecting localization cloud quantification limit

The results presented in this article (Figure 2(B)) for determining the localization cloud identification limit are based on experimentally measured localization uncertainty, molecular dissociation rate, and the search region used in mean-shift clustering. Here, the influence of these parameters on the quantification limit—specifically the maximum localization cloud density $\sigma_{\text{cloud}}^{\text{max}}$ identifiable with at least 90% accuracy—is systematically evaluated.

As shown in Figure S2, the molecular dissociation rate $k_{\text{off}}$, which influences the average number of localizations per localization cloud, has negligible impact on $\sigma_{\text{cloud}}^{\text{max}}$. In contrast, reducing localization uncertainty (i.e., decreasing the standard deviation of the Gaussian distribution from which localization coordinates are sampled) increases $\sigma_{\text{cloud}}^{\text{max}}$. This indicates that, for a given spatial distribution, a greater number of localization clouds can be reliably identified when localizations are determined with higher precision. This result is expected, as better localization precision reduces the spatial overlap between nearby localization clouds. However, the benefit of improved localization precision is less pronounced in clustered molecular distributions. Even with high localization accuracy, $\sigma_{\text{cloud}}^{\text{max}}$ increases only marginally under strong clustering, suggesting that spatial overlap remains a limiting factor in such systems. As previously shown in,[S2] an optimal search region exists for mean-shift clustering. Varying the search region for data generated with 8 nm localization uncertainty confirms this: $\sigma_{\text{cloud}}^{\text{max}}$ initially increases with search region size but decreases beyond an optimal point, due to merging of distinct localization clouds.

In experimental settings, the search region is often selected empirically through visual inspection of clustering results. Alternatively, it can be guided by the observed localization uncertainty, with a typical rule of thumb being twice the maximum measured uncertainty.

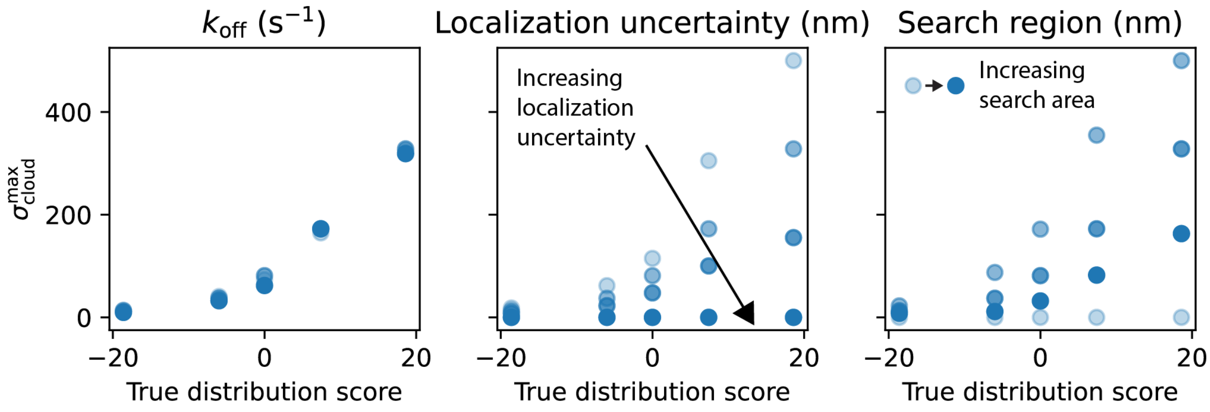


Figure S2: Effects of molecular dissociation rate $k_{off}$, localization uncertainty and mean-shift clustering search region on the localization cloud identification limit.

#### S3 Areas of interpretability in quantified distribution scores

In Figure 3, regions of interpretability are identified—combinations of true surface molecular density $\sigma_{mol}$ and molecular sampling ratio $s_r$ for which quantified spatial distributions can be confidently interpreted. These regions vary with the underlying true spatial distribution. Figures S3 and S4 illustrate how the size and shape of these interpretability regions evolve across different spatial distributions. Notably, clustered systems exhibit substantially smaller interpretability areas compared to random and dispersed systems. As the true distribution approaches complete spatial randomness (i.e., distribution score near zero), the area of interpretability narrows, indicating that fewer combinations of $\sigma_{mol}$ and $s_r$ yield statistically interpretable distribution scores.

The significantly smaller interpretability regions observed for clustered systems, compared to random and dispersed systems, highlight a key limitation of PAINT in quantifying spatial distributions under clustered molecular arrangements. While this may not always be a critical drawback, particularly when alternative molecular targets such as linker molecules provide more interpretable spatial information, the challenge becomes more pronounced when a high-density, clustered system is the object of interest. In such cases, expanding the interpretability region is essential for reliable distribution quantification.

One approach to achieve this is by increasing the spatial resolution of PAINT, for instance through the use of RESI (Resolution Enhancement by Sequential Imaging).[S7] The impact of enhanced PAINT resolution is shown in Figure S5. Here, localization coordinates are sampled from a Gaussian distribution with a standard deviation of 0.8 nm, tenfold lower than the 8 nm uncertainty used throughout this study. To ensure consistency, the mean-shift clustering search region is also reduced proportionally to 3 nm.

As shown in Figure S5, the area of interpretable quantified distribution scores expands considerably for a system with comparable clustering (true distribution score = -18.63), allowing a wider range of $\sigma_{\mathrm{mol}}$ and $s_{\mathrm{r}}$ combinations to yield statistically interpretable results. However, it is important to note that experimental methods enabling such high resolution often suffer from poor scalability with molecular density and may require prolonged acquisition times (ranging from hours to days) to collect sufficient statistics for robust quantification. This underscores the fundamental trade-off between imaging parameters and surface molecular complexity in achieving high-throughput and precise quantification in PAINT.

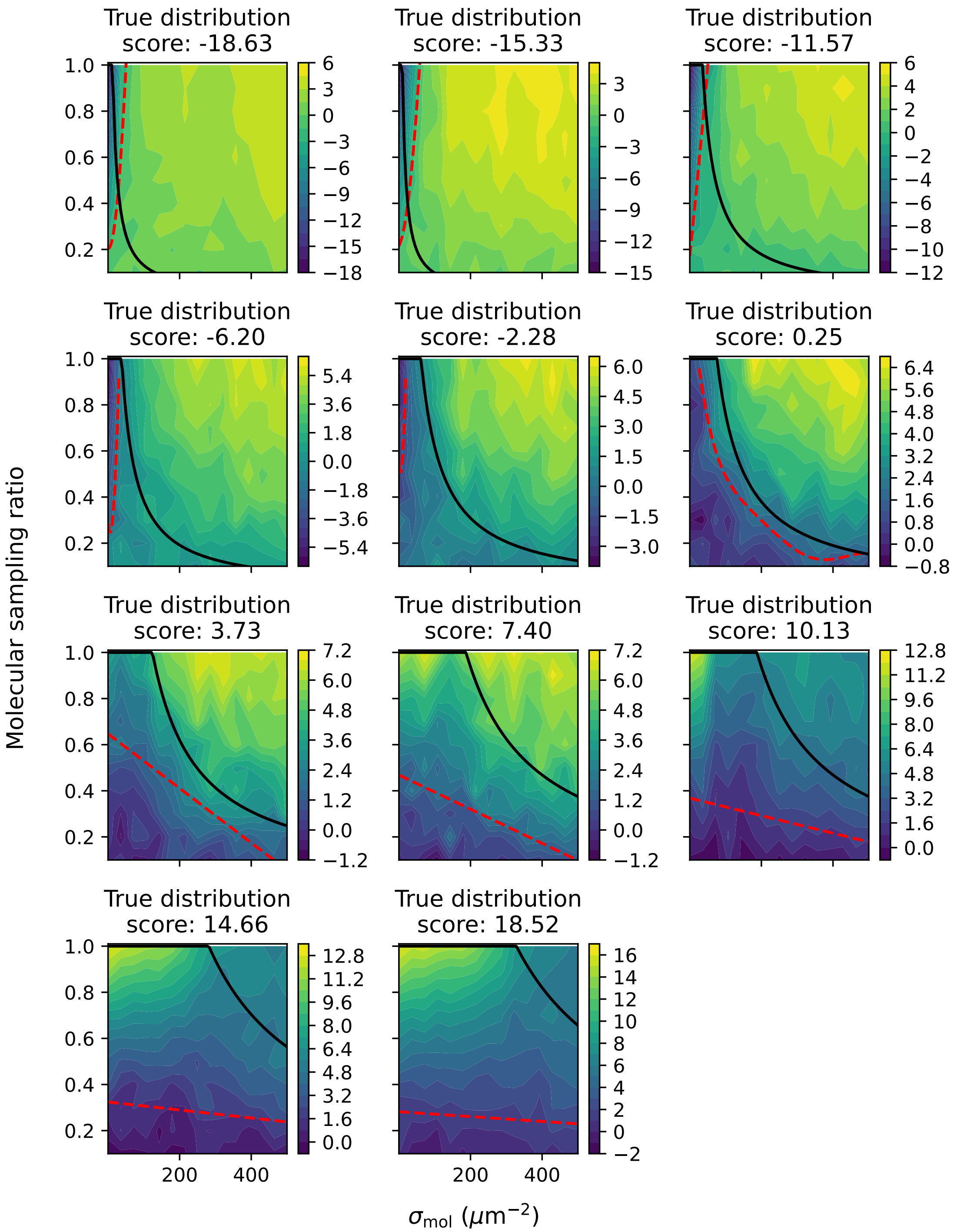


Figure S3: Contour map of observed distribution score as a function of true molecular density $\sigma_{\mathrm{mol}}$, molecular sampling ratio $s_{\mathrm{r}}$, and true spatial distribution. The black line indicates the maximum allowable sampling ratio, defined by the localization cloud identification limit. Red dotted lines mark the distribution score thresholds corresponding to the 5% significance level, distinguishing significantly clustered or dispersed distributions from complete spatial randomness. Distribution score less than -$z_{0.05}$ (-1.65) indicates clustering, while distribution score more than $z_{0.05}$ indicates dispersity.

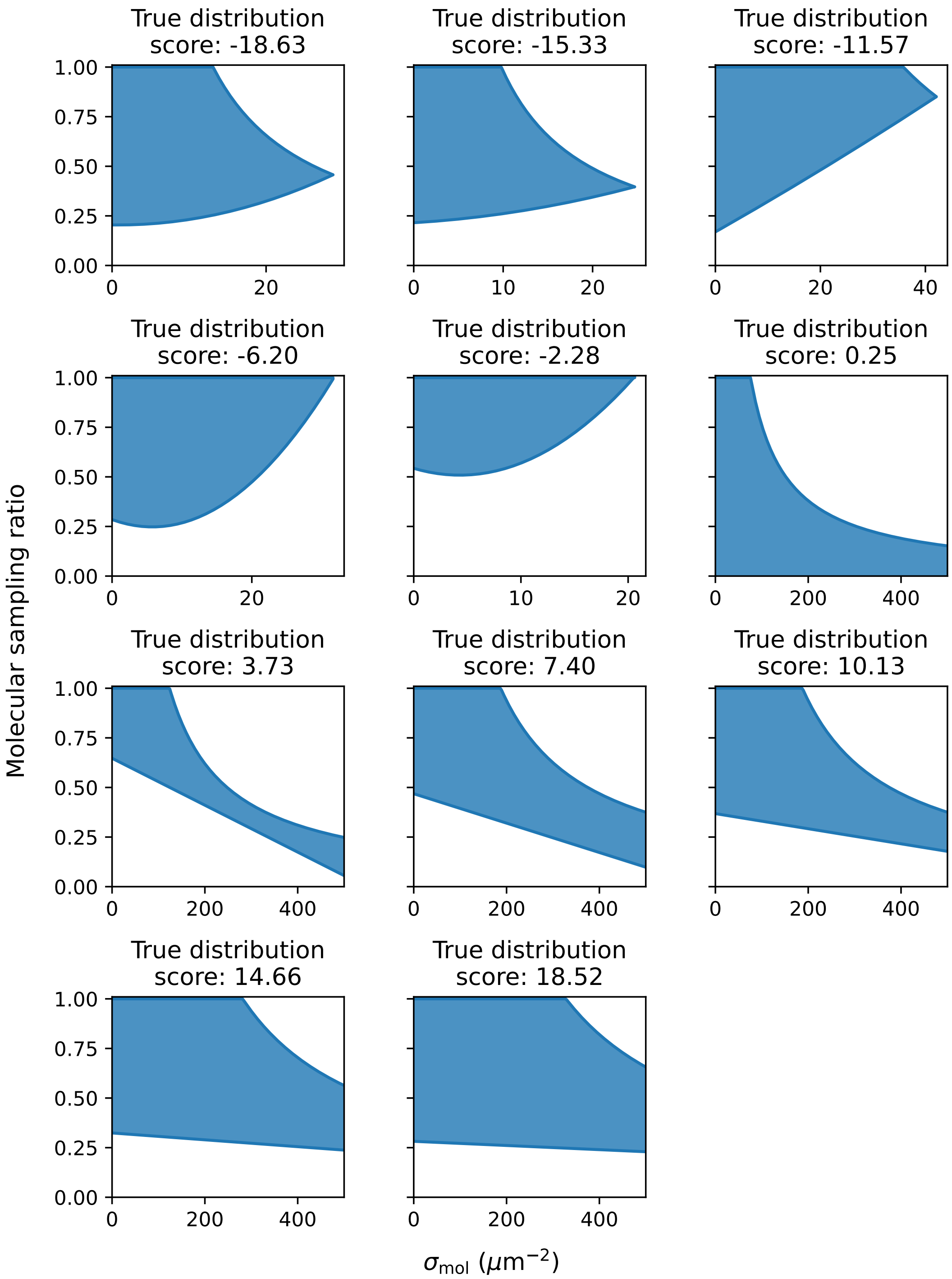


Figure S4: Area of interpretable quantified distribution scores as a function of true molecular density $\sigma_{mol}$ and molecular sampling ratio $s_r$, across varying true spatial distributions. The shaded regions indicate combinations of $\sigma_{mol}$ and $s_r$ for which the observed distribution score can be confidently interpreted, based on statistical significance and localization cloud identification accuracy. Distribution score less than $-z_{0.05}$ (-1.65) indicates clustering, while distribution score more than $z_{0.05}$ indicates dispersity. Note: the x-axis scales differ between plots.

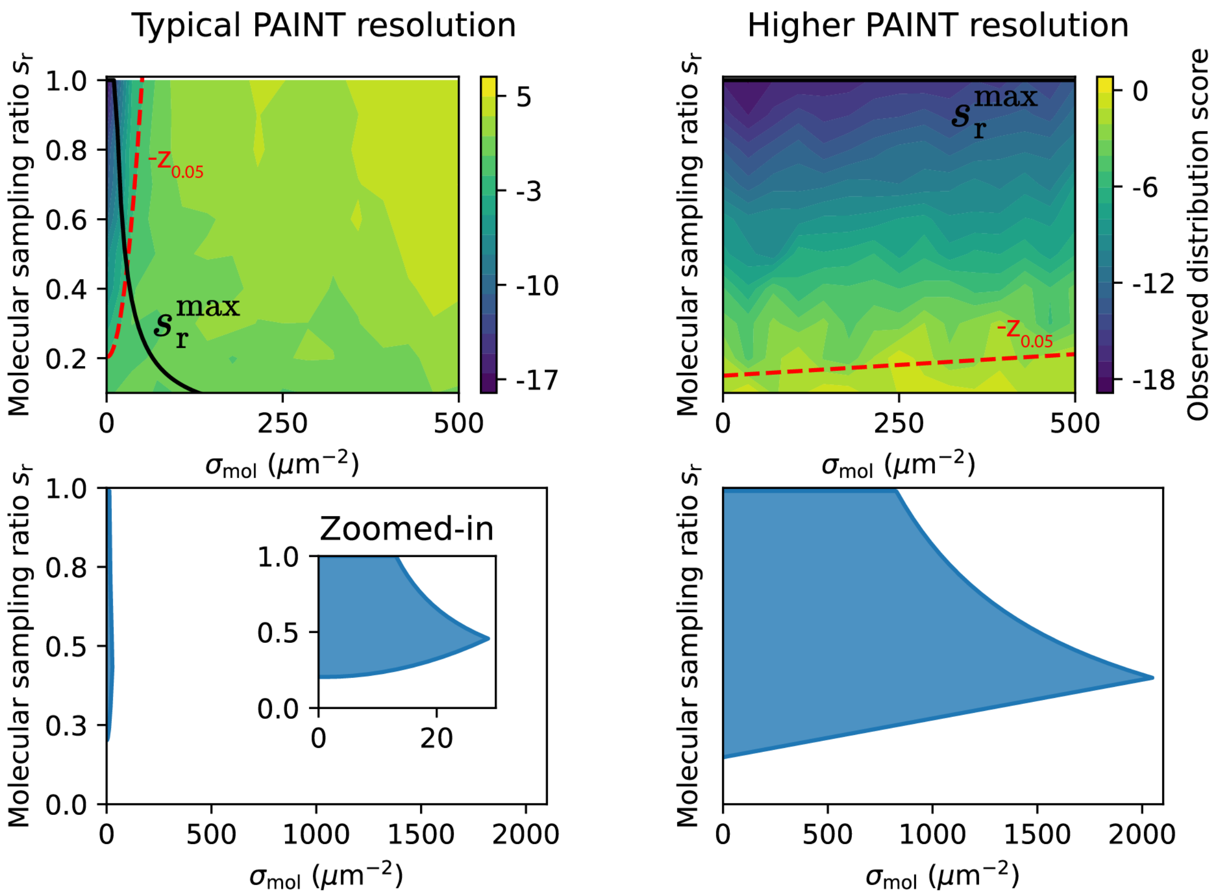


Figure S5: Contour maps of observed distribution scores and corresponding interpretability regions as functions of true molecular density $\sigma_{\mathrm{mol}}$ and molecular sampling ratio $s_{\mathrm{r}}$ for a clustered system (true distribution score = -18.63). For the typical PAINT resolution case, synthetic localizations are generated with a localization precision of 8 nm and identified using a mean-shift clustering search region of 30 nm. For the high-resolution case, localizations are generated with a precision of 0.8 nm and identified using a 3 nm search region. Shaded regions indicate combinations of $\sigma_{\mathrm{mol}}$ and $s_{\mathrm{r}}$ where the observed distribution score is statistically interpretable.

#### S4 Probe scores

##### S4.1 Range of molecular density quantifiable with at least 90% accuracy per probe

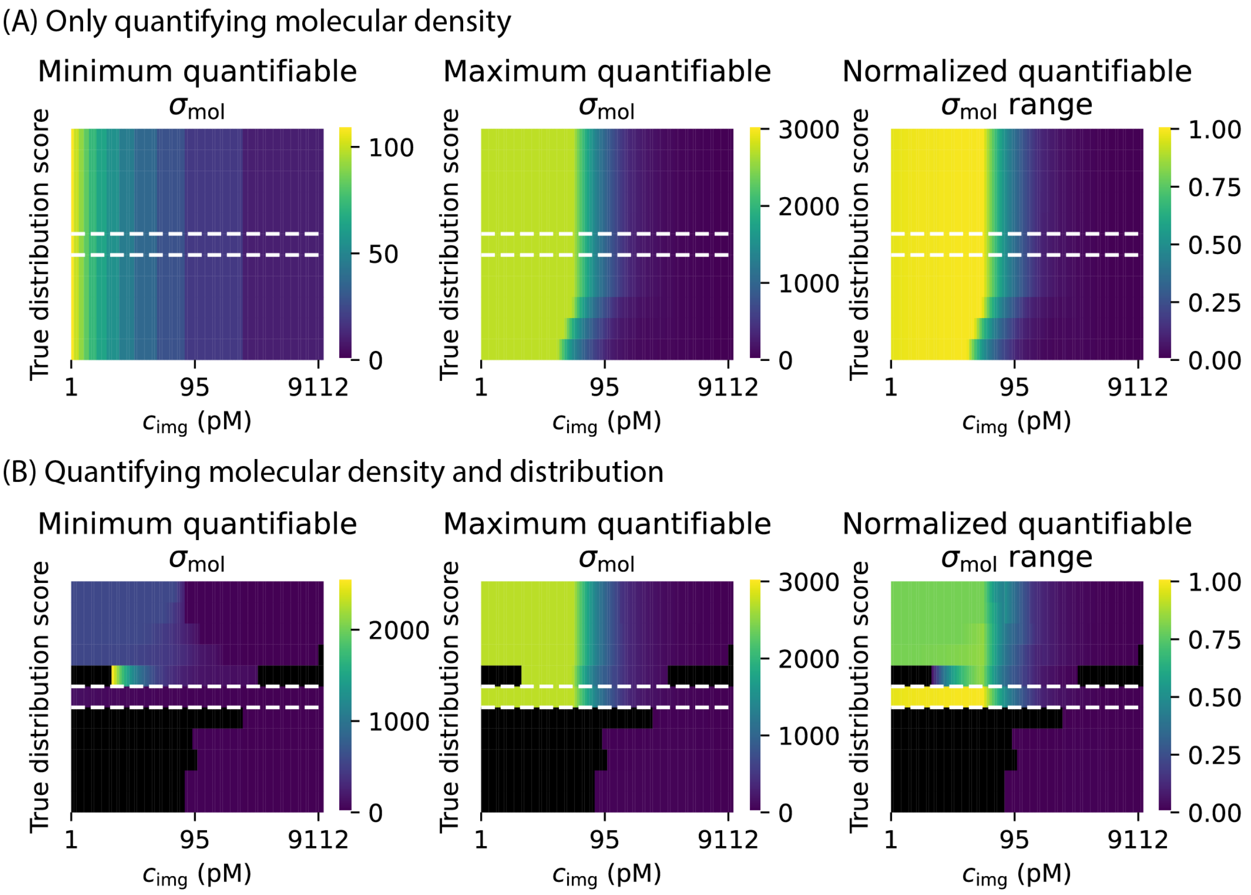


Figure S6: Maps of minimum and maximum quantifiable molecular density $\sigma_{\text{mol}}$, and the corresponding normalized quantifiable density range, for different combinations of true spatial distribution and probe concentration $c_{\text{img}}$. (A) Quantification based on density accuracy limits only. (B) Quantification based on both density accuracy limits and interpretability of distribution scores.

Figure S6 shows the range of molecular densities $\sigma_{\text{mol}}$ that can be quantified with at least 90% accuracy and interpretable distribution scores for a DNA probe with reference kinetics of $k_{\text{on}} = 10^6$ M$^{-1}$ s$^{-1}$ and $k_{\text{off}} = 1$ s$^{-1}$. For each probe concentration $c_{\text{img}}$ and true spatial distribution, the minimum and maximum quantifiable $\sigma_{\text{mol}}$ values are determined based on the quantification limits and interpretability criteria established in Figures 2 and 3. The resulting density range is then normalized by the total simulated $\sigma_{\mathrm{mol}}$ range to produce a map of the normalized quantifiable density range. This map is used throughout the study to evaluate the impact of quantification constraints and probe kinetics, and serves as the basis for calculating probe scores.

#### S4.2 Calculation of probe scores

A probe score is assigned to each probe based on its kinetic constants and is derived from the map of quantifiable density range. When considering only the density quantification limit, the normalized quantifiable density range is first averaged across all true spatial distributions for each probe concentration $c_{\mathrm{img}}$. The probe score is then defined as the maximum of these mean values. When both density and spatial distribution quantification are considered, the map is divided into three spatial regimes, namely clustered, random, and dispersed, highlighted in red in Figure S7. For each regime, the mean normalized quantifiable density range is calculated across $c_{\mathrm{img}}$, and the maximum value is taken as the probe score.

This definition reflects the objective of identifying probes that support reliable quantification over a broad range of true molecular densities $\sigma_{\mathrm{mol}}$ and spatial distributions, regardless of the probe concentration used. A high probe score therefore indicates strong and versatile quantification performance across varying experimental and molecular conditions.

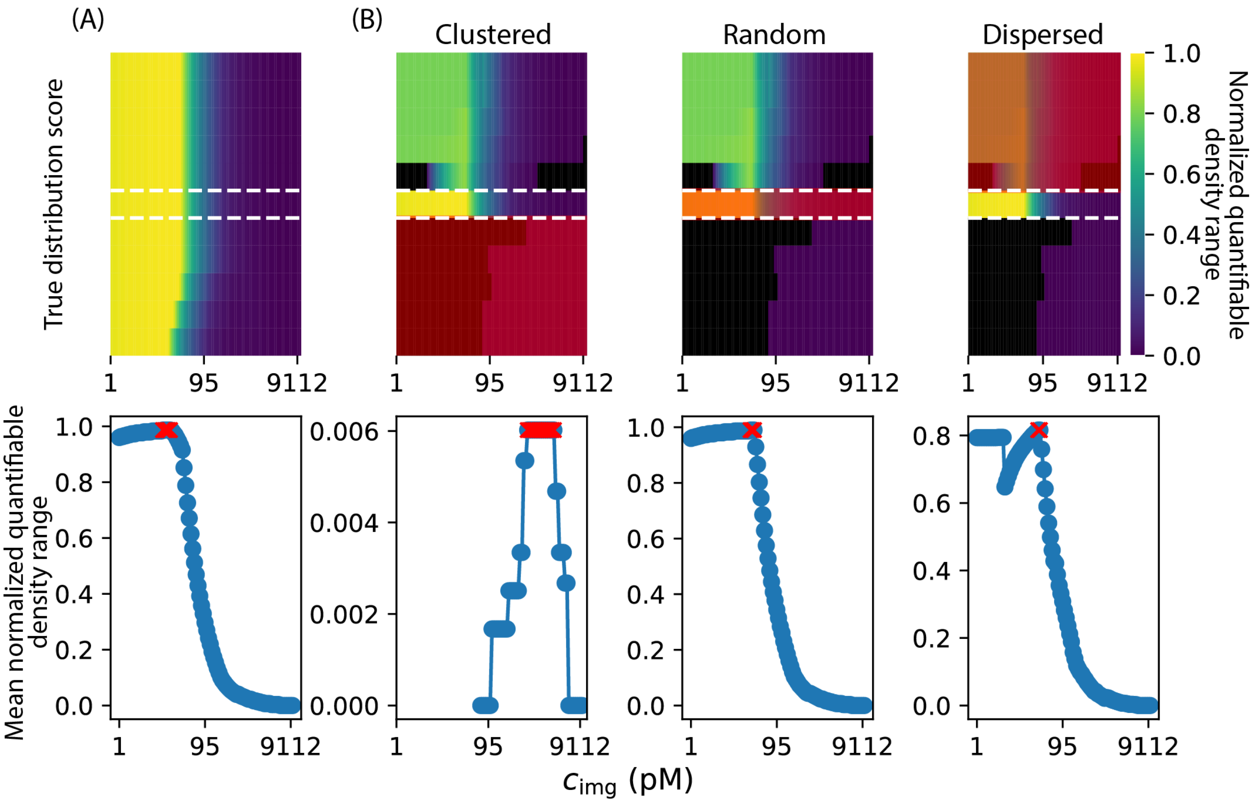


Figure S7: Maps of normalized quantifiable density range (top row) and corresponding mean normalized quantifiable density range across varying probe concentrations (bottom row). Probe scores are defined as the maximum of the mean values. (A) Quantification based on density accuracy limits only. (B) Quantification based on both density accuracy limits and interpretability of distribution scores.

#### S4.3 Effect of quantification limit and probe kinetics

In this work, four quantification limits or criteria are proposed to ensure molecular quantification with at least 90% accuracy, as summarized in Table S7. These limits impact the map of normalized quantifiable density range differently depending on the probe kinetic constants. Figure S8 illustrates how these limits sequentially affect the quantifiable density range for four representative sets of probe kinetics.

The $\sigma_{\mathrm{PSF}}^{\mathrm{max}}$ limit constrains the range of probe concentrations $c_{\mathrm{img}}$ that support high-accuracy quantification, particularly for probes with low $k_{\mathrm{off}}$ or high $k_{\mathrm{on}}$. For example, a probe with $k_{\mathrm{on}} = 10^4$ M$^{-1}$ s$^{-1}$ and $k_{\mathrm{off}} = 1$ s$^{-1}$ enables a broad quantifiable density range across a wide $c_{\mathrm{img}}$ interval. While the $\sigma_{\mathrm{cloud}}^{\mathrm{max}}$ limit has only a minor effect on the density map within the simulated ranges of $c_{\mathrm{img}}$ and acquisition time, the $\sigma_{\mathrm{event}}^{\mathrm{min}}$ limit, which accounts for the statistical requirement to overcome molecular undersampling, can substantially affect the quantifiable range depending on probe kinetics. Probes with lower $k_{\mathrm{on}}$ fail to generate sufficient event statistics at low $c_{\mathrm{img}}$, leading to regions with no quantifiable density (indicated by black squares). One particularly interesting case is the probe with $k_{\mathrm{on}} = 10^6$ M$^{-1}$ s$^{-1}$ and $k_{\mathrm{off}} = 0.1$ s$^{-1}$. Although a high $c_{\mathrm{img}}$ would typically support precise quantification, this probe is strongly limited by the $\sigma_{\mathrm{PSF}}^{\mathrm{max}}$ constraint. Furthermore, very low densities at this probe concentration do not yield sufficient event statistics, meaning that no quantifiable $\sigma_{\mathrm{mol}}$ exists within this regime.

When the interpretability of spatial distribution is also considered, the quantifiable density map is further constrained by the required sampling ratio $s_{\mathrm{r}} = \sigma_{\mathrm{cloud}}/\sigma_{\mathrm{mol}}$. The map is divided into three spatial regimes: clustered, random, and dispersed. For the random case, the interpretability area aligns closely with the $\sigma_{\mathrm{cloud}}^{\mathrm{max}}$ constraint, and thus does not significantly alter the trend. For dispersed systems, one might expect only minor impact due to the broad interpretability region. However, in the four kinetic examples shown, certain regions remain non-quantifiable because the minimum required $s_{\mathrm{r}}$ is not met. In this case, the sampling ratio is dictated by the statistical threshold necessary to classify the distribution as dispersed (i.e., distribution score $\geq z_{0.05}$). For clustered systems, quantifiable densities are restricted to very low values due to the inherently narrow interpretability region. Notably, the probe concentration range that allows precise quantification also varies significantly with probe kinetics.

Figures S9 and S10 further illustrate how probe kinetics influence the map of quantifiable density range under two scenarios: considering only density accuracy limits, and considering both density and distribution interpretability. Although the nine probes shown in these figures have similar overall probe scores, the $c_{\text{img}}$ regimes over which high-accuracy quantification is achievable differ markedly. In some cases, the effective concentration window becomes narrower, broader, or shifts depending on kinetic parameters. This emphasizes the importance of tailoring imaging parameters to the specific kinetic characteristics of the PAINT probe in use.

Table S7: Summary of quantification limit defined in this work.

| Quantification limit | | Notes |
|---|---|---|
| 1 | $\sigma_{\text{PSF}}^{\text{max}}$ limit | Avoid PSF overlap |
| 2 | $\sigma_{\text{cloud}}^{\text{max}}$ limit | Avoid localization cloud overlap |
| 3 | $\sigma_{\text{event}}^{\text{min}}$ limit | Ensure sufficient statistics |
| 4 | Area of interpretability | Accurate estimates of molecular position and statistical thresholding |

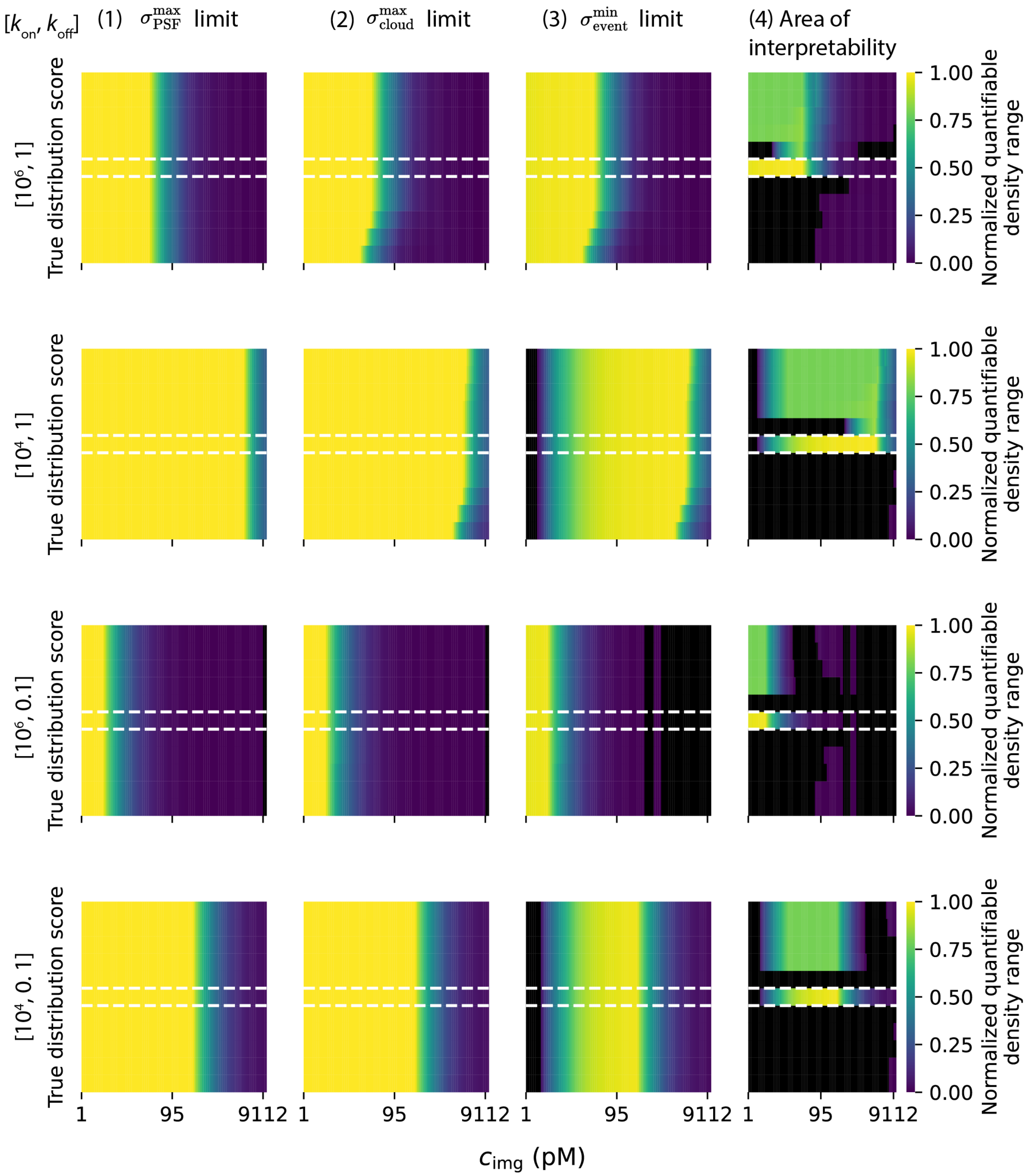


Figure S8: Evolution of the map of quantifiable density range as quantification limits are applied sequentially. Four representative probe kinetics are shown to illustrate how each limit, namely $\sigma_{PSF}^{max}$, $\sigma_{cloud}^{max}$, $\sigma_{event}^{min}$, and the area of interpretability, progressively constrains the range of molecular densities $\sigma_{mol}$ that can be quantified with at least 90% accuracy.

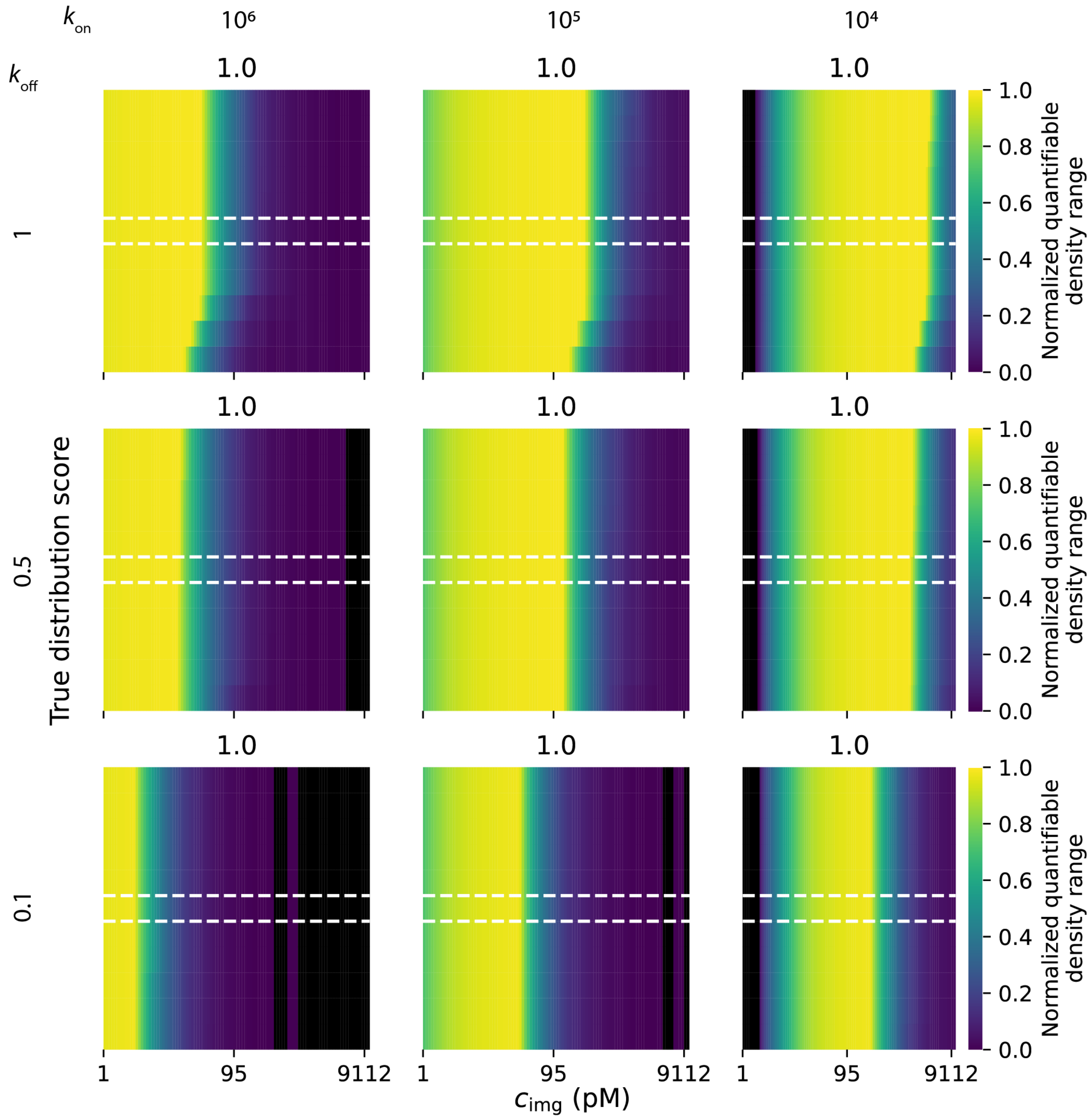


Figure S9: Effect of probe kinetics on the map of quantifiable density range when only density accuracy is considered. The figure illustrates how different combinations of probe association and dissociation rates influence the range of molecular densities $\sigma_{\mathrm{mol}}$ that can be quantified with at least 90% accuracy. Each subplot is titled with the corresponding probe score.

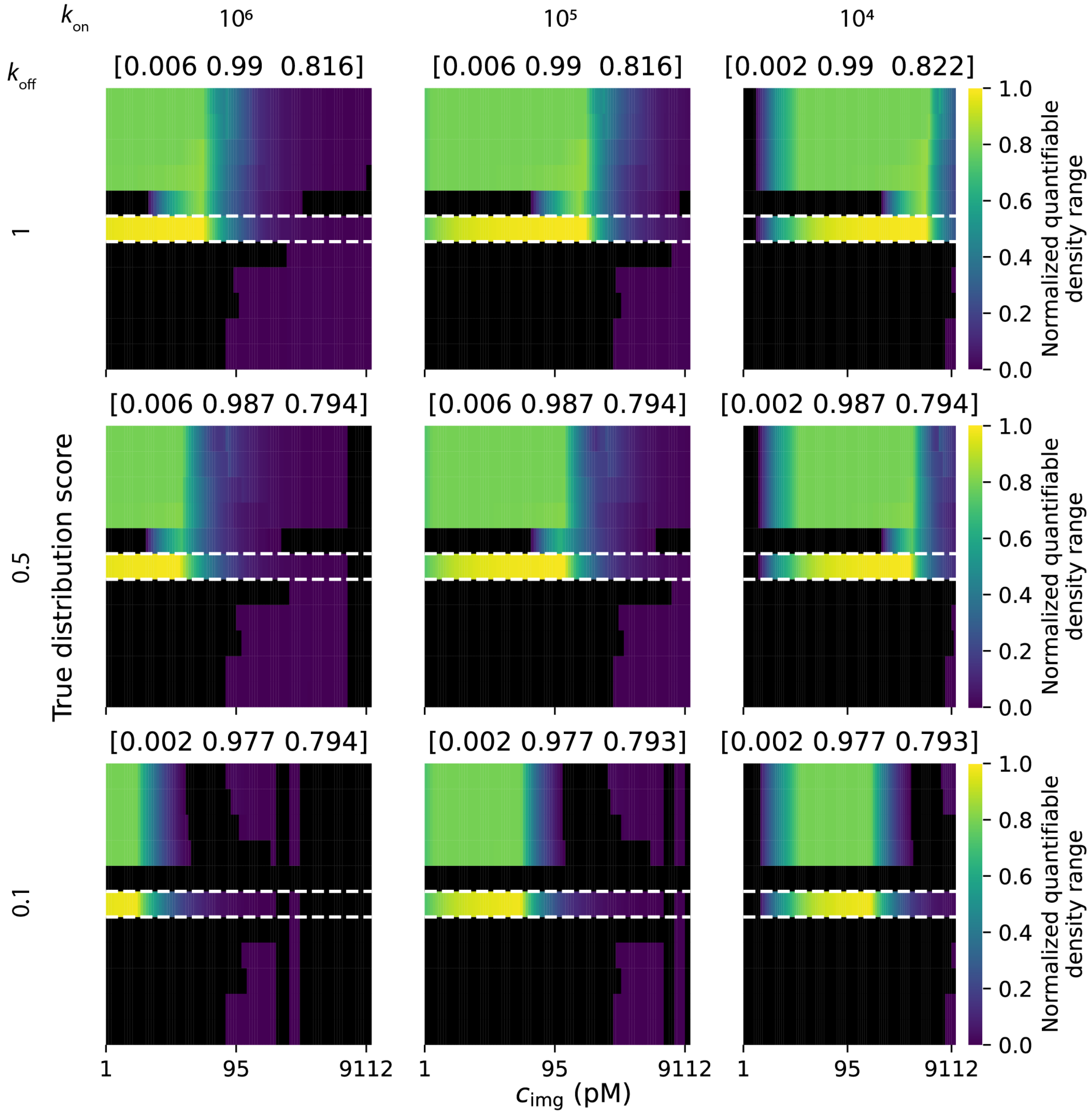


Figure S10: Effect of probe kinetics on the map of quantifiable density range when density accuracy and distribution interpretability is considered. The figure illustrates how different combinations of probe association and dissociation rates influence the range of molecular densities $\sigma_{mol}$ that can be quantified with at least 90% accuracy. Each subplot is titled with the corresponding probe score, reported for clustered, random, and dispersed distributions.

#### S4.4 Effect of PAINT resolution

As shown in Figure S5, the area of interpretability expands significantly when higher PAINT resolution is employed, as in the case of RESI. This expansion directly increases the normalized quantifiable density range achievable for a given probe. Figure S11 illustrates the effect of enhanced PAINT resolution across different probe kinetics, exemplified for a true spatial distribution score of $-18$. Notably, higher resolution leads to a substantial increase in quantifiable density range, up to approximately 60-fold for probes with high dissociation rates ($k_{\mathrm{off}}$) across the simulated $c_{\mathrm{img}}$ range. Even for probes with lower $k_{\mathrm{off}}$ values, the gain remains significant, with an increase of roughly 20-fold. In the Figure, the maximum normalized quantifiable density range is computed for each probe across the tested $c_{\mathrm{img}}$ range. The results show a modest expansion in the range of kinetic parameters that support high-accuracy quantification under clustered conditions. In particular, probes with higher $k_{\mathrm{off}}$ benefit more from increased resolution, enabling slightly broader quantification capability. These findings reinforce that higher PAINT resolution can alleviate some of the limitations associated with quantifying high-density, clustered molecular distributions.

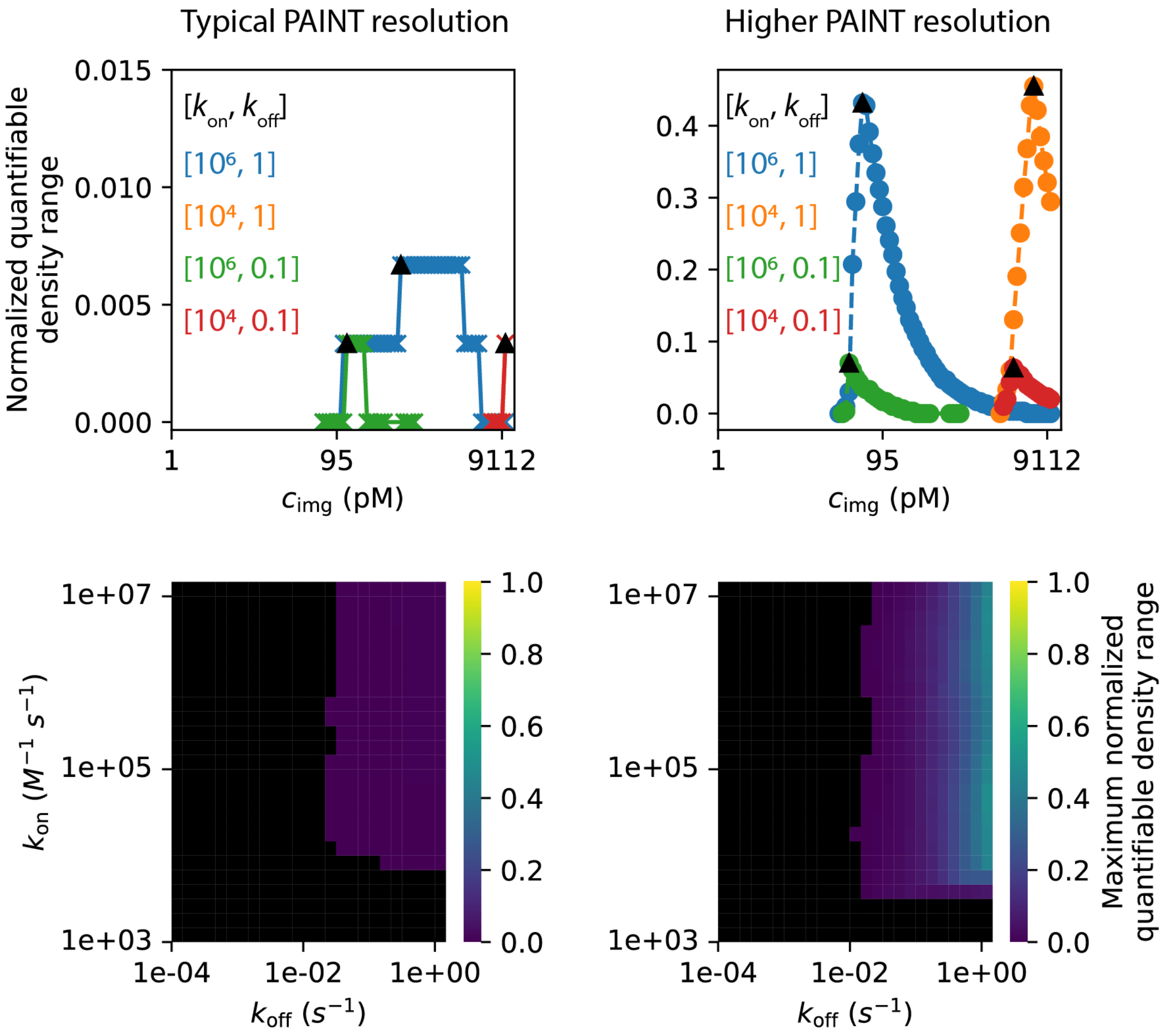


Figure S11: Effect of PAINT resolution on the quantifiable density range for clustered systems (true distribution score = -18). (Top row) Normalized quantifiable density ranges are shown for four example probe kinetic constants, comparing typical PAINT resolution and higher-resolution conditions. Black triangles indicate the maximum normalized quantifiable density range for each probe. (Bottom row) Maps of the maximum normalized quantifiable density range across varying probe kinetic constants, derived from quantification limits based on typical and higher PAINT resolutions.

### S5 Additional information

#### S5.1 Experimental details for NN experimental validation dataset

##### S5.1.1 Materials and chemicals

Glass coverslips ($22 \times 40$ mm, thickness #1.5, Epredia) were obtained from VWR. Custom-made flow cell stickers with an approximate internal volume of 20 $\mu$L were obtained from Grace Biolabs (USA). Poly(l-lysine)-grafted poly(ethylene glycol) (PLL-g-PEG) with a grafting ratio of 3.5 was purchased from SuSoS (Switzerland). The molecular weight of the PLL backbone and PEG side chains are 20 and 2 kDa respectively. Azide functionalized PLL-g-PEG (Nanosoft Biotechnology LLC, USA) is composed of a 15 kDa PLL backbone and 2 kDa PEG chain with a grafting ratio of 5. Fluorescent nanoparticles (0.2 $\mu$m, yellow-green, Molecular Probes) were used as fiducial markers. PBS tablets, NaCl, and $Mg_2Cl$ were purchased from Sigma-Aldrich, and Tween 20, EDTA, and Tris-HCl were purchased from Merck Life Science. The ssDNA oligonucleotides (standard desalting and HPLC purification for chemically modified DNA) were purchased from IDT (Integrated DNA Technologies). PBS buffer was prepared by dissolving 1 tablet of PBS in 200 mL of MilliQ water. 1 M of NaCl dissolved in PBS was used as the high-salt (HS) buffer in this study. The imaging buffer (Buffer B) consists of 10 mM $Mg_2Cl$, 5 mM Tris-HCL, 1 mM EDTA and 0.05% Tween 20.

Table S8 shows the ssDNA molecules that were used in this work. Sequence #1 is pre-hybridized with Sequence #2, and conjugated to the low-fouling polymer to prepare the ssDNA-functionalized surface. For the protein-functionalized surface, Sequence #3 is first conjugated to the low-fouling polymer surface, followed by the hybridization of the ssDNA-conjugated protein (Sequence #4). Sequence #5 is the ssDNA imager strand used for DNA-PAINT experiment. Sequence #3 and #4 are kindly provided by collaborators and the conjugation of ssDNA molecules to the protein is described in.[S8]

Table S8: ssDNA sequences used in this work.

| # | 5' end group | Sequence | 3' end group |
|---|---|---|---|
| 1 | None | CGATTCGAGAACGTGACTGCTTTTT | DBCO |
| 2 | None | GCAGTCACGTTCTCGAATCGAACATTATTACA | None |
| 3 | DBCO | GTGCGGCAGGGGTAAGACCA | None |
| 4 | Protein | TTTTTTTTGGTCTTACCCCTGCCGCAC | None |
| 5 | ATTO647N | TTGTAATAATG | None |

##### S5.1.2 Coverslip surface functionalization

The surface preparation procedure follows from.[S2] Briefly, the coverslips were washed by 10 min of sonication in isopropanol and MilliQ baths respectively. After the sonication steps, the coverslips were dried under nitrogen flow and placed under 1 min of oxygen plasma to oxidize the surface. A flow cell sticker was then attached to the coverslip and a polymer mixture solution was immediately added to the flow cell and incubated for approximately 3 hours. The polymer mixture solution consists of 1% v/v PLL-g-PEG-N3/PLL-g-PEG, for ssDNA-functionalized surface, or 10% v/v PLL-g-PEG-N3/PLL-g-PEG, for protein-functionalized surface, at a final combined concentration of 0.5 mg/mL in MilliQ. After the polymer incubation step, the solution in the flow cell was then aspirated to remove unbound or loosely bound polymer molecules and replaced with solution containing 2 $\mu$M of partially double-stranded DNA (for ssDNA-functionalized surface) or 2 $\mu$M of ssDNA linker (for protein-functionalized surface) in HS buffer. The DNA-containing solution was incubated for 3 days. For the protein-functionalized surface, the surface was flushed with 100 $\mu$L of HS buffer before replacing the buffer solution with 75 nM of protein-ssDNA conjugate molecules in HS buffer. The solution was incubated for 10 min prior to PAINT imaging preparation.

##### S5.1.3 PAINT imaging and analysis

Fiducial markers were prepared by first sonicating the stock solution for 5 minutes to disaggregate clusters of the particles. After sonication, the stock suspension was diluted 10000 times using PBS or HS buffer and the diluted suspension was subjected to 5 min sonication.

Before flowing in the fiducial markers, the flow cells were flushed with 100$\mu$L of PBS or HS buffer. Then, the buffer in the flow chamber was exchanged with the suspension of fiducial markers, and the fiducial markers were allowed to sediment and attach to the substrate for 5 minutes. After this procedure, the unattached fiducial markers were flushed away with 100$\mu$L of Buffer B or HS buffer, and 25 pM of ssDNA probe solution (in Buffer B) or 1 nM of Fab probe solution (in HS buffer) was added into the flow chamber. For the PAINT experiment on the ssDNA-functionalized surface, combination of PBS and Buffer B was used. On the other hand, only HS buffer was used during the PAINT imaging preparation for the protein-functionalized surface. PAINT imaging was performed on Oxford Nanoimager with a TIRF configuration. Fluorescence was recorded using a 100×, 1.4 NA oil immersion objective, passed through a beam splitter to obtain a green and a red channel. Images were acquired with an exposure time of 100 ms under 640 nm laser and 0.05 mW of 532 nm laser illumination simultaneously for 1 hour. The laser power for the 640 nm laser was 12 mW and 31 mW for the ssDNA probe and Fab probe respectively. The raw images were then analyzed with ThunderSTORM,[S1] an open-source plug-in in ImageJ, to extract the localizations of the fluorescence emissions. Analysis of the localizations obtained was described in.[S2] Briefly, the localizations were filtered based on their properties and split into $5 \times 5$ $\mu$m$^2$ region of interest ROI. Then, the spatio-temporal information of the localization data per ROI was used to obtain the binding event density. The binding event density was used as inputs for the BiSC-based analysis and NN-based analysis to estimate true surface molecular density. The p-values provided in this work are obtained via Welch's t-test statistic.

#### S5.2 Sobol analysis results

Table S9 presents the results of the Sobol sensitivity analysis. For each input and output parameter pair, the first-order index ($S_i$), total-order index ($S_t$), and the interaction contribution ($S_t - S_i$) are reported. The low values of interaction contributions (i.e., $S_t - S_i < 0.1$) indicate minimal interaction effects between input parameters, suggesting that the variability in PAINT outputs is predominantly driven by the individual input parameters themselves.

Table S9: Sobol sensitivity analysis results for PAINT output metrics. First-order indices ($S_i$), total-order indices ($S_t$), and interaction contributions ($S_t - S_i$) are shown for each input parameter. The values quantify the relative influence of individual parameters and their interactions on the variability of PAINT outputs.

| Input parameter | $S_i$ | $S_t$ | $S_t - S_i$ |
|---|---|---|---|
| Output parameter: $\sigma_{\mathrm{PSF}}$ | | | |
| $k_{\mathrm{on}}$ | 0.361 | 0.367 | 0.005 |
| $\sigma_{\mathrm{mol}}$ | 0.136 | 0.136 | 0.001 |
| $k_{\mathrm{off}}$ | 0.272 | 0.278 | 0.006 |
| $c_{\mathrm{img}}$ | 0.221 | 0.225 | 0.004 |
| $t$ | 0.001 | 0.005 | 0.003 |
| Output parameter: $\sigma_{\mathrm{cloud}}$ | | | |
| $k_{\mathrm{on}}$ | 0.482 | 0.493 | 0.011 |
| $\sigma_{\mathrm{mol}}$ | 0.204 | 0.205 | 0.001 |
| $k_{\mathrm{off}}$ | 0.002 | 0.003 | 0.001 |
| $c_{\mathrm{img}}$ | 0.271 | 0.282 | 0.010 |
| $t$ | 0.030 | 0.032 | 0.002 |
| Output parameter: $\sigma_{\mathrm{event}}$ | | | |
| $k_{\mathrm{on}}$ | 0.492 | 0.499 | 0.007 |
| $\sigma_{\mathrm{mol}}$ | 0.185 | 0.185 | 0 |
| $k_{\mathrm{off}}$ | 0 | 0.004 | 0.004 |
| $c_{\mathrm{img}}$ | 0.283 | 0.289 | 0.006 |
| $t$ | 0.032 | 0.033 | 0.001 |
